# Tetracycline-Regulated Inducible CB_2_ Expression in AtT20 Cells: A Functional Assay for Quantifying Ligand Efficacy

**DOI:** 10.64898/2026.02.26.708391

**Authors:** Tahira Foyzun, Mark Connor, Humayra Zaman, Michael Kassiou, Annukka Kallinen, Marina Junqueira Santiago

**Author notes:** **Corresponding author**: Mark Connor. Macquarie Medical School, Faculty of Medicine, Health and Human Sciences, Macquarie University, NSW 2109, Australia.

## Abstract

**Introduction:** Cannabinoid receptor-2 (CB_2_) is an emerging therapeutic target for chronic and inflammatory pain, cancer, and neurological disorders. Understanding the efficacy of CB_2_ ligands is crucial for future drug design and development.

**Aims:** We aimed to establish a simple and robust system to control CB_2_ expression using a tetracycline-regulated mammalian expression system (T-REx), to enable application of the Black and Leff operational model to measure the operational efficacy (τ) of CB_2_ ligands.

**Methods:** Ligand-induced hyperpolarisation of AtT20 cells transfected with T-REx and human CB_2_ was measured by FLIPR membrane potential assay. Maximal and submaximal responses of the CB_2_ ligands were produced by regulating CB_2_ expression using tetracycline. Data were fitted to the operational model of receptor depletion to quantify the efficacy of seven ligands. Additionally, the maximal initial rate of signalling (IR_max_), another putative measure of ligand efficacy, was determined.

**Results:** AK-F-064, CP55940 and 2-AG exhibited similar efficacy with a τ values of 11.4, 11 and 10.4 respectively, while anandamide (AEA) had the lowest efficacy (τ=1.07) among the tested agonists. The rank order of operational efficacy and IR_max_ was similar and was estimated as: AK-F-064 = CP55940 = 2-AG > 5F-AB-PICA = WIN55212-2 > HU-308 = AEA.

**Conclusion:** This inducible expression system provides a reliable platform for quantifying and comparing CB_2_ ligand efficacy using the operational model. This approach may facilitate more precise CB_2_-targeted drug development and can be readily extended to other GPCR targets.

## Introduction

Cannabinoid receptors-2 (CB_2_) [1] are predominantly expressed in immune cells and peripheral tissues, in contrast to the more extensively studied CB_1_ receptor, which is highly expressed in the central nervous system. CB_2_ has been the focus of more attention as research in preclinical models suggests it being a viable target for novel therapeutic strategies in alleviating symptoms in autoimmune diseases, inflammation, and pain [1–4]. Unlike CB_1_, which also regulates pain and inflammation, activation of CB_2_ does not produce obvious psychotropic and psychiatric effects, consistent with its very limited expression in the central nervous system [5–7]. Despite several compounds reaching clinical trial stage, no agonist that selectively targets CB_2_ reached the market [8].

Efficacy - a measure of the ability of a drug to activate its target - is an important property of a drug that plays a crucial role in its clinical effectiveness [9–11]. However, the relative efficacy of CB_2_ ligands is understudied. Many pharmacological approaches to measure agonist efficacy rely on the measurement of binding affinity and functional concentration-response relationships. These approaches often overlook the dynamics of receptor activation in a physiological membrane environment, as well as the presence of spare receptors, also known as ‘receptor reserve’ (those are in excess to produce the maximal effect when occupied by an agonist) [12–16]. Without considering spare receptors, it is impossible to predict the proportional occupancy required to achieve a maximal response, and these assays cannot quantify or differentiate the relative efficacy of agonists, particularly those capable of eliciting maximal responses.

To consider the influence of spare receptors, while quantifying efficacy, Black and Leff developed the operational model of agonism, which subsumed the spare receptors along with the parameters of intrinsic efficacy and proposed a unique parameter, τ’ - the ‘transducer ratio’ or ‘operational efficacy’ of an agonist [17]. One way to apply the operational model is to fit two concentration-response curves, using conditions where maximum and submaximal responses of an agonist can be measured. This is usually achieved by receptor depletion in a system, where a fraction of the available receptors is partially inactivated using alkylating agents or an irreversible antagonist [17–19]. This technique was successfully applied to determine the CB_1_ agonist efficacy using an irreversible antagonist, AM6544 [20], however, the lack of a suitable irreversible antagonist for CB_2_ limits the use of this method.

Therefore, this study aimed to develop an alternative strategy using T-REx (Tetracycline-Regulated Mammalian Expression) to alter CB_2_ density and facilitate measurement of agonist efficacy. This approach controls the receptor expression on the cell surface rather than inactivating them. Using tetracycline to regulate the binding of the Tet-repressor protein to a site upstream of the CB_2_ receptor gene, CB_2_ receptor expression could be controlled to allow an agonist to elicit both maximum and submaximal responses, which facilitated the use of the operational model to fit the data and provide the key parameters (K_A_ and E_max_) required to calculate efficacy. In the present study, we developed a stable cell line where CB_2_ expression was controlled by an inducible expression system, T-REx, to establish a new method for assessing the efficacy of a range of CB_2_ agonists, using a robust high-throughput FLIPR (Fluorescence Imaging Plate Reader) membrane potential assay.

## Materials and Methods

### Chemicals and reagents

The sources of synthetic CB_2_ agonists and somatostatin were provided in Supplementary Table 1. AK-F-064 and 5F-AB-PICA were a gift from Michael Kassiou (University of Sydney). Antibiotics blasticidin, hygromycin B gold, and zeocin were from InvivoGen. Vectors pOG44, FRT/lacZeo2, and pcDNA6/TR were purchased from Thermo Fisher Scientific and pcDNA5/FRT/TO with the CB_2_ construct was from GenScript. Cell culture reagents Dulbecco’s Modified Eagle Medium (DMEM), Leibovitz’s medium (L-15), Fetal Bovine Serum (FBS), Phosphate-Buffered Saline (PBS), trypsin-EDTA solution, and Penicillin-Streptomycin (P/S) were obtained from Gibco. Bovine Serum Albumin (BSA) and Tetracycline Hydrochloride were purchased from Merck, Australia. Transfection reagent FuGENE®HD was from Promega.

### Generation of inducible AtT20Flp-InT-REx-CB_2_ cell lines

To achieve the controlled expression of CB_2_ in our study, we used the T-REx system (Supplementary Figure 1), developed by Yao and colleagues and commercialized by Invitrogen [21]. The wild-type AtT20 cells (ATCC: CRL-1795) were first stably transfected with the pFRT/lacZeo2 vector that integrates an FRT site into the transfected cells to generate zeocin-resistant cells. A zeocin-resistant clone was isolated and propagated to generate the AtT20Flp-In host cell line [22].

Subsequently, the pcDNA6/TR vector, which expresses the tetracycline repressor (TR) protein and the blasticidin resistance gene, was introduced into AtT20Flp-In host cells to generate the clones of AtT20Flp-InT-REx host cell line. The transfected clones were isolated following blasticidin selection. Seven individual clones were randomly selected and screened for a functional response to 100 nM of somatostatin in the FLIPR assay (described below). The clones that exhibited similar responses to somatostatin as untransfected AtT20Flp-In cells were selected for further transfection.

Finally, the expression vector pcDNA5/FRT/TO, carrying the human CB_2_ with an N-terminus triple-HA tag and the Flp recombinase expression plasmid, pOG44, were co-transfected into the selected AtT20Flp-InT-REx clones (in a 1:9 ratio) using FuGENE®HD (FuGENE: DNA=4:1) (Supplementary Table 2). The pOG44 facilitates the integration of the CB_2_-containing vector into the genome through the FRT site, generating stable AtT20Flp-InT-REx-CB_2_ cells. An Empty vector (EV) control cell line was generated by inserting the pcDNA5/FRT/TO vector (without the gene of interest) into AtT20Flp-InT-REx clones following a similar procedure. Transfected cells resistant to 150 μg/mL of hygromycin B and 10 μg/mL of blasticidin over 5 passages were selected.

### Cell culture

AtT20Flp-InT-REx-CB_2_ and EV were propagated in 75 cm^2^ tissue culture flasks (Corning) using DMEM supplemented with 10% FBS, 1% P/S, hygromycin (80 μg/mL), and blasticidin (5 μg/mL). The cells were incubated at 37°C in an atmosphere with 5% CO_2_ humidified air. In this study, the passage number of transfected cells ranged between 5 and 25.

### FLIPR Membrane potential assay

We used a proprietary membrane potential assay (MPA) to measure the changes in cellular membrane potential, triggered by CB_2_-mediated activation of the endogenously expressed G protein-gated inwardly rectifying potassium (GIRK) channels in AtT20 cells [23,24]. This assay was conducted using the FLIPR assay kit (blue) (Molecular Devices, USA), as detailed by Knapman et al [25]. The dye was diluted to 50% of the manufacturer’s recommended concentrations with a low-potassium Hank’s Balanced Salt Solution (HBSS; composition in Supplementary Table 3).

For each assay, 80-90% confluent cells were detached with trypsin-EDTA solution, followed by centrifugation and resuspension in L-15 medium containing 1% FBS, 1% P/S, and 15 mM glucose. The cells were plated in black-walled, clear-bottomed 96-well microplates (Corning) at a volume of 90 µL/well. Resuspended cells with or without tetracycline were used to assess efficacy following 16 hours of incubation at 37 °C in humidified ambient air. Dye was added to the cells, and the plate was transferred to the FlexStation 3 microplate reader (Molecular Devices) operated by SoftMax Pro 7 software. The emission and excitation of the dye were continuously measured at wavelengths of 565 and 530 nm, respectively.

The drugs were prepared at 10x the desired final concentration in well by serial dilution in HBSS supplemented with 0.01% BSA and 1% DMSO. Drugs were added to the cells after a 120s-baseline reading, with responses measured every 2 seconds for a total run time up to 5 minutes unless otherwise noted. HBSS+ 0.01% BSA+1% DMSO was used as a control (final in well 0.001% BSA and 0.1% DMSO). The experiments were carried out in duplicate at least five to seven times for each compound, and the average of these duplicates was used for analysis.

### Data analysis

The changes in membrane potential because of agonist-induced GIRK channel activation was expressed as the maximum percent change in fluorescence compared to baseline after subtraction of any changes produced by the addition of control. All the raw data from SoftMax Pro were analysed in Excel and then fitted into GraphPad Prism 10.

### a. Fitting into the Operational model

#### Operational model of receptor depletion

The operational efficacy tau (τ) and functional affinity (K_A_) of each agonist were estimated by fitting the maximal and submaximal response data of the individual agonist into the Black and Leff operational model of receptor depletion (five parameters-Basal, *K*_A_, *Effect*_max_, τ, and transducer slope) in GraphPad Prism. The equation is presented as:

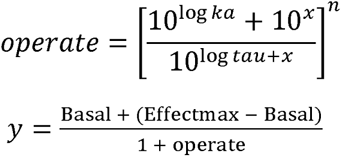

The basal and transducer slope (n) was constrained to zero and 1, respectively. The highest *Effect*_max_ was calculated from the agonists’ CRCs using a three-parameter nonlinear regression equation (with Basal constrained to zero) in Prism and shared among the agonists tested on that day.

#### Operational model of Partial agonists

Some tested agonists had small responses under low-receptor conditions, and the data could not be reliably fitted into the operational model because of the lack of a robust CRC. Hence, we determined the efficacy of these agonists relative to the reference high efficacy agonist CP55940 and fitted the data into the extended form of the operational model called the “Operational model of Partial agonist” in GraphPad Prism [26]. The dose-response data for both CP55940 and the tested agonists obtained from the tetracycline-induced cells were analysed by fitting them into the following equation:

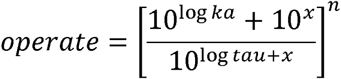

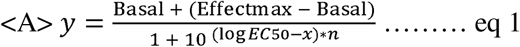

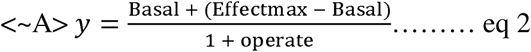

Equation 1, indicated by <A>, applies only to the dataset obtained from the reference agonist and represents a variable-slope dose–response fit. The second line, indicated by <∼A>, applies to all tested agonists except the reference and incorporates the operational model to assess the partial agonists’ affinity and efficacy. The E_max_ of the system was estimated from CP55940 data, which then allowed operational fitting of the tested partial agonists and determination of their affinity and efficacy.

#### Four-parameter nonlinear regression equation

For the somatostatin dose-response curve, the data were fitted into the four-parameter nonlinear regression equation to obtain the concentration-response curve (CRC).

### b. Fitting into the Pharmechanics model: Measurement of kinetics of ligand efficacy

Quantifying the signal generation rate of agonist-bound receptors as the initial rate (IR_max_) of signalling is a newly described approach for assessing agonist efficacy. We analysed the time-course data of CB_2_-mediated membrane hyperpolarisation using the kinetic analysis method outlined by Hoare (2020) and quantified ligand efficacy in kinetic terms [27]. The response of the maximally effective concentration of five agonists was measured in uninduced AtT20Flp-InT-REx-CB_2_ cells with low receptor expression. The low receptor density ensures the absence of spare receptors. The time course data of signal generation of agonist-occupied CB_2_ obtained from FLIPR assay were fitted into the user-defined equation: [Pharmechanics] “Baseline then fall and rise to steady state” time course equation of the Graph-Pad Prism 10, which is:

Y= IF (X<X0, Baseline, Baseline+(-SteadyState) *(1-D*exp(-K_1_*(X-X0)) +(D-1) *exp(-K_2_*(X-X0))))

Here, Y = signal level,

SteadyState = ligand-specific steady-state response

D= fitting constant,

K_1_ and K_2_ = the rate constants of the fall and rise phase, respectively

X = time, X0 = signal start time after the application of agonist

Then the IR_max_ (initial rate of signalling of the agonist-bound receptor) and trough response (the maximum response below the baseline of the receptor signal generation) were determined in PRISM using the following equations:

Initial rate, IR_max_ = SteadyState × (Dk_1_ – (D – 1) k_2_)

Trough Response= SteadyState × (1-D × exp (-K_1_/ (k_2_-K_1_) × In ((D-1) × K_2_/ (D× K_1_))) + (D-1) × e

#### Statistical analysis

The data are presented as the mean ± SEM from 5-7 independent experiments (unless otherwise stated), conducted in duplicate. Statistical analyses were performed using GraphPad Prism 10. D’Agostino and Pearson’s method was used to assess the normality and lognormality of the data. The statistical comparison of the operational efficacy and initial rate of the agonists was assessed through one-way ANOVA followed by Tukey’s test. An unpaired two-tailed t-test was performed to determine significant differences in maximal response between tetracycline-induced and uninduced cells.

## Results

### Optimisation of inducible expression of CB_2_

We initially aimed to identify optimal tetracycline concentrations that ensured a reliable difference in the Emax of the high-efficacy agonist CP55940 in AtT20Flp-InT-REx-CB_2_ cells. This was necessary to apply the operational model to a situation where one condition (high receptor expression) defined the maximal system response to receptor activation, and the other condition (low receptor expression) had no spare receptors for CP55940. To achieve this, cells were incubated with 0.03, 0.1, and 1 µg/mL of tetracycline (Figure 1). The maximum response of CP55940 elicited by 0.1 and 1 µg/mL tetracycline induction was similar (29.2 ± 1.9% and 24.4 ± 1.7%, respectively, p = 0.2074). Therefore, we chose to use 0.1 µg/mL tetracycline to limit potential adverse effects of exposure to higher tetracycline concentrations. Induction with 0.03 µg/mL tetracycline produced a lower response (E_max_= 20.3 ± 1.8%) compared to 0.1 µg/mL induction (p = 0.0092). The expression system was leaky with substantial basal expression, sufficient for CP55940 to elicit the submaximal responses (E_max_=15.3 ±1.4%) without tetracycline induction. This response was significantly lower than that observed following 0.1 µg/mL tetracycline induction (p <0.0001). Thus, the submaximal responses of agonists were measured in uninduced cells.

**Figure 1:**
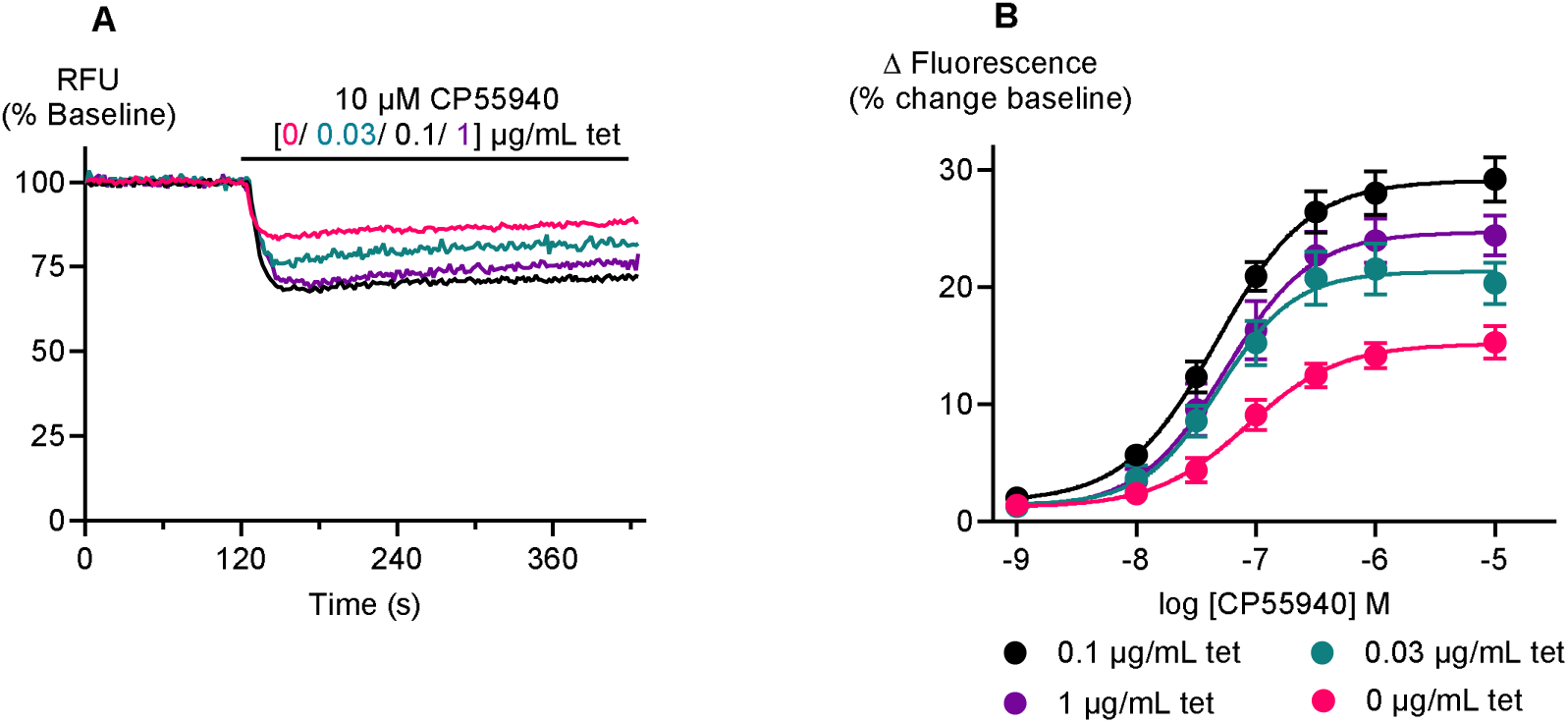
Optimisation of tetracycline concentration.

The dose-response curve of CP55940 in AtT20Flp-InT-REx-CB_2_ cells incubated with different concentrations of tetracycline (tet). (A) Representative traces are examples of the changes in fluorescence signal (relative fluorescence units normalised to baseline fluorescence) in tetracycline-induced (with 0.03, 0.1, and 1 µg/mL of tetracycline) and uninduced cells in the FLIPR assay after the addition of CP55940. (B) Concentration-response curves for CP55940 under uninduced and induced (with 0.03, 0.1, and 1 µg/mL of tetracycline) conditions. Each point represents the mean ± SEM of 5-7 independent determinations.

### Functional Validation of the T-REx system

To investigate whether there was any potential impact of transfection and tetracycline induction on GIRK channel signalling, we assessed the responses of native somatostatin receptors (SSTR) in AtT20Flp-InT-REx wild type (WT), EV, and CB_2_ cells [28,29]. The activation of the SSTR by somatostatin (SST) was measured in the FLIPR assay. The E_max_ (38 ± 1, 37.0 ± 1.1, and 37.4 ± 0.5%) and pEC_50_ (9.5 ± 0.1, 9.3 ± 0.07 and 9.6 ± 0.05) values of SST across the tested cell lines (Figure 2A and 2B) were not different (p = 0.8279 and 0.1523 for E_max_ and pEC_50_, respectively).

**Figure 2:**
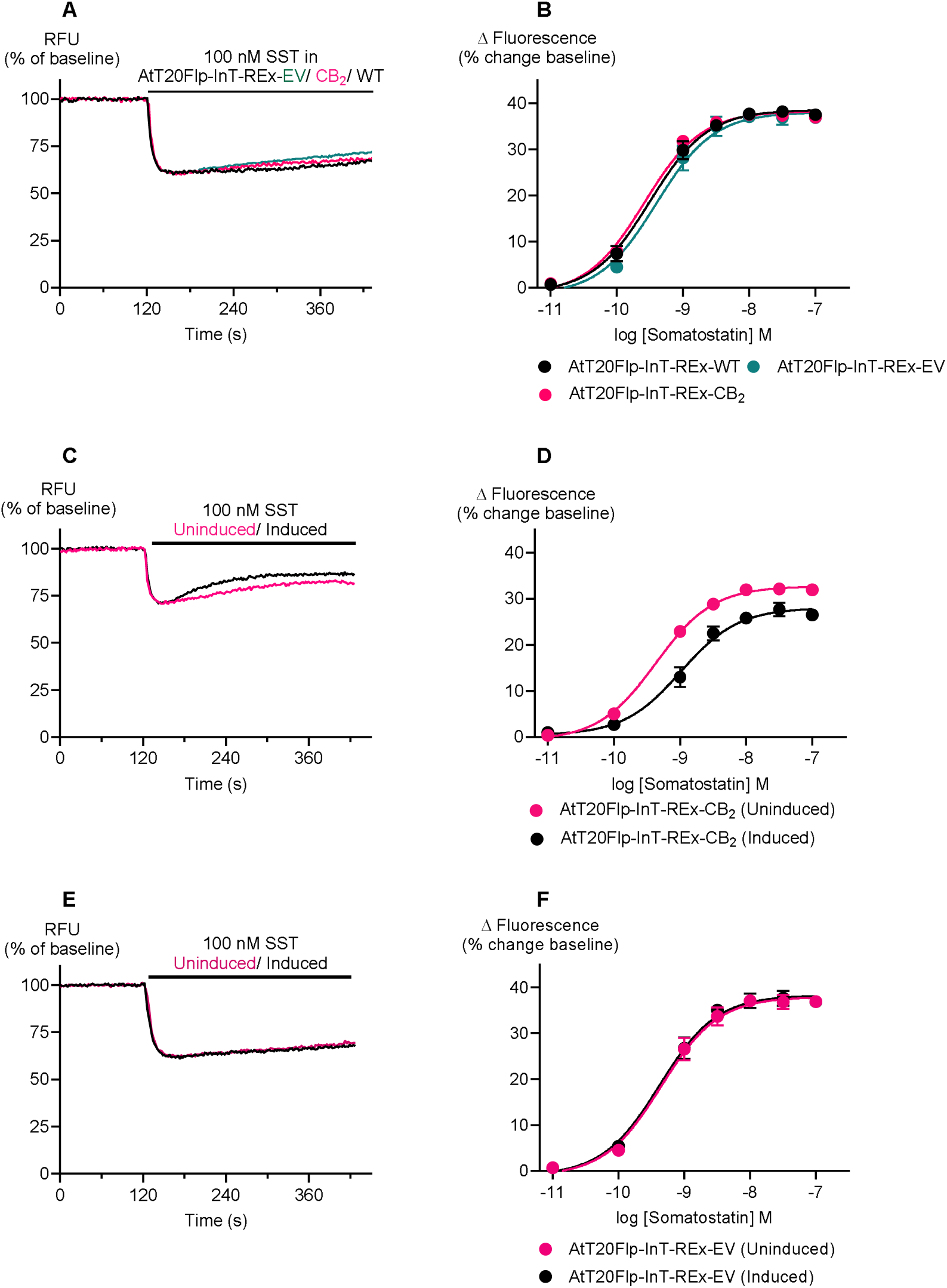
Validation of the T-REx system.v

Notably, SST responses were different between the uninduced and induced (with 0.1 µg/mL tetracycline) CB2-transfected cells. The SST exhibited higher E_max_ and pEC_50_ values (32.2± 0.9% and 9.3 ± 0.07) in uninduced cells compared to their induced counterparts (26.5 ± 1.1% and 8.9 ± 0.06) (p = 0.0012 and 0.0024 for E_max_ and pEC_50_, respectively) (Figure 2C and 2D). However, no significant differences were observed in the SST response in EV-transfected cells with or without tetracycline, which rules out the possibility of tetracycline interference with the SSTR or the underlying signalling pathways (Figure 2E and 2F). Furthermore, there was no significant difference in the E_max_ of the GIRK channel activator ML-297 in AtT20Flp-InT-REx-CB_2_ cells with (0.1 µg/mL) and without tetracycline incubation (p = 0.9745). This implies that the GIRK channel expression was preserved during the transfection process (Supplementary Figure 2), but that expression of higher levels of CB_2_ receptors altered the expression or functional coupling of SST receptors [30].

Somatostatin (SST) responses were assessed by measuring responses at endogenous somatostatin receptors (SSTR) in AtT20Flp-InT-REx cells. The traces are examples of the changes in fluorescence signal (relative fluorescence units normalised to baseline fluorescence) in the FLIPR assay after the addition of SST. Panels 2A and B show example traces and CRC for SST at uninduced AtT20Flp-InT-REx-wild type, AtT20Flp-InT-REx-EV and AtT20Flp-InT-REx-CB_2_ cells, confirming that transfection did not alter the native SSTR signalling. Panels 2C and 2D, and 2E and 2F show the SST response in AtT20Flp-InT-REx-CB_2_ and AtT20Flp-InT-REx-EV cells, respectively, incubated with 0.1µg/mL of tetracycline and without tetracycline. Tetracycline alone did not affect SSTR signalling. CRCs data are shown as mean ± SEM (n = 5).

### Determination of the operational efficacy (**τ**) and functional affinity (pK_A_) of CB_2_ agonists using the T-REx system

In this study, we assessed seven CB_2_ ligands: the commonly used reference agonists CP55940 and WIN55212-2, the synthetic cannabinoid 5F-AB-PICA, the experimental PET (positron emission tomography) ligand AK-F-064 [31], the highly CB_2_-selective synthetic agonist HU-308 and two endocannabinoids, 2-arachidonoyl glycerol (2-AG) and anandamide (AEA). The operational efficacy (τ) and functional affinity (K_A_) of all ligands were measured by fitting the data to the operational model of receptor depletion, except for HU-308 and AEA. Both HU-308 and AEA showed low responses under uninduced conditions; their responses were instead measured under induced conditions, compared with CP55940 and analysed using the operational model of partial agonism. CRCs for all seven drugs tested are shown in Figures 3, 4 and 5, efficacy and pK_A_ are presented in Table 1, and E_max_ and EC_50_ values are available in Supplementary Table 4.

**Figure 3:**
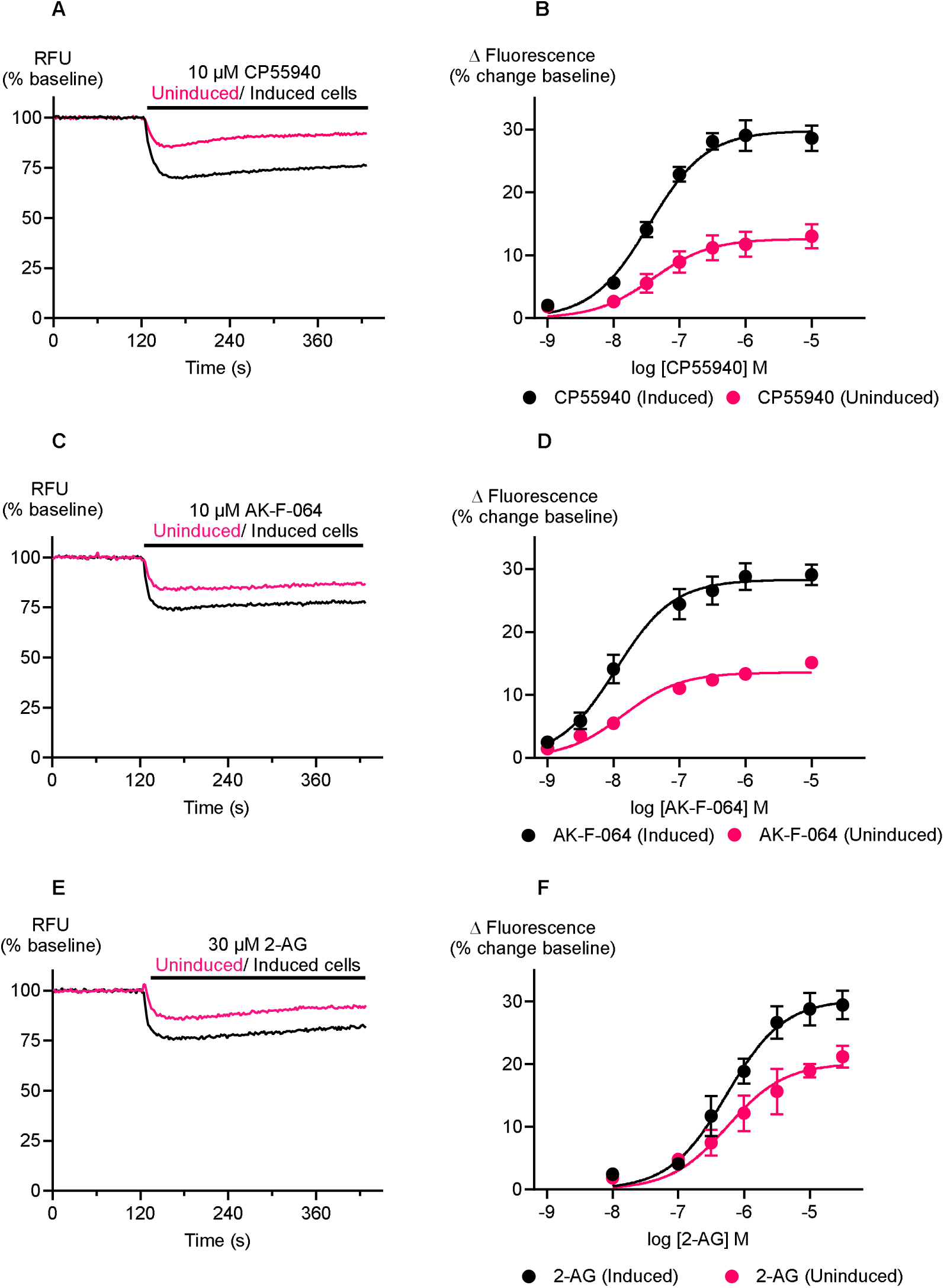
Signalling of high-efficacy CB_2_ agonists in AtT20Flp-InT-REx-CB_2_ cells incubated with or without tetracycline, in FLIPR assay.

**Figure 4:**
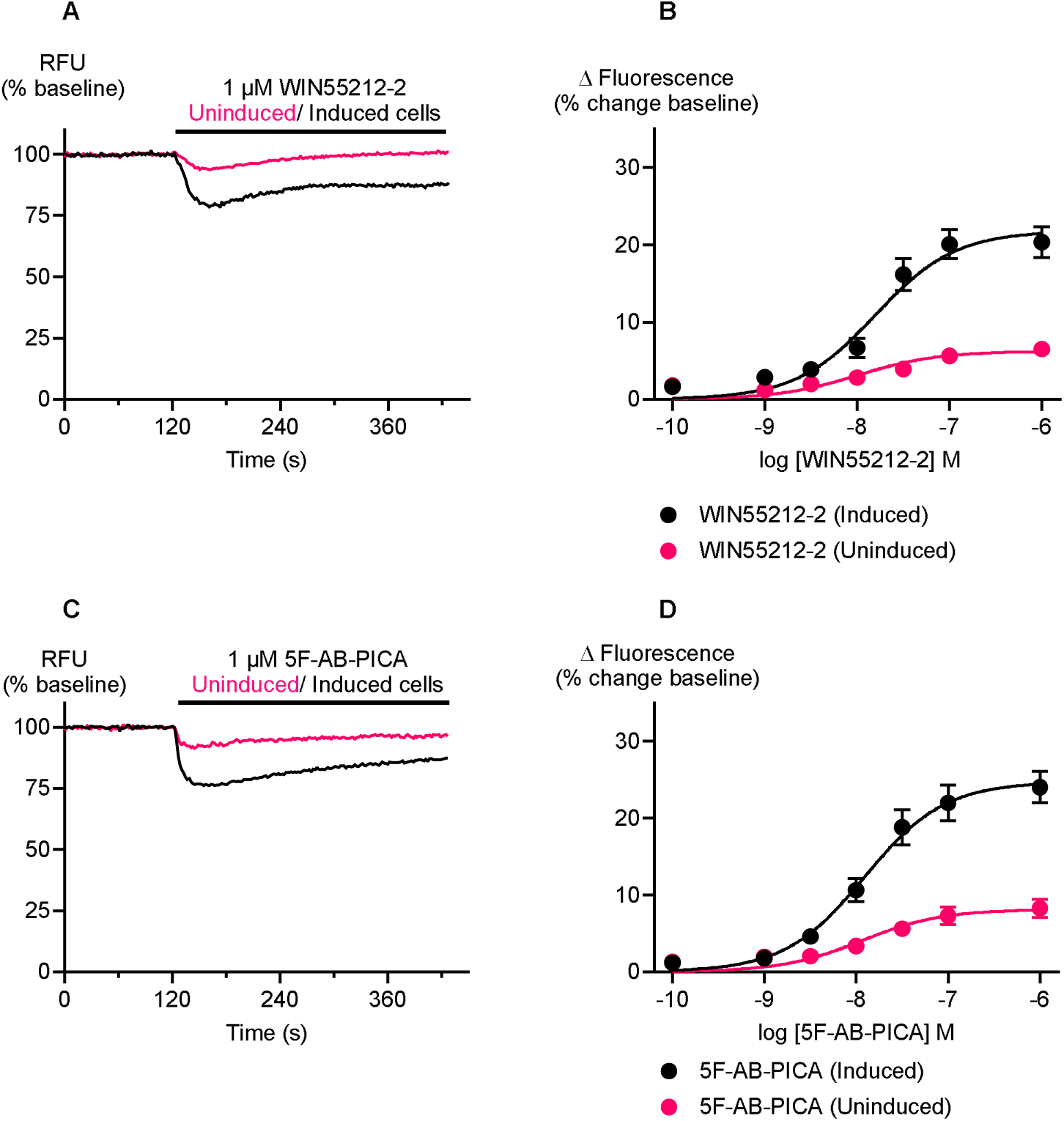
Signalling of moderate efficacy CB_2_ agonists in AtT20Flp-InT-REx-CB_2_ cells incubated with or without tetracycline, in FLIPR assay.

**Figure 5:**
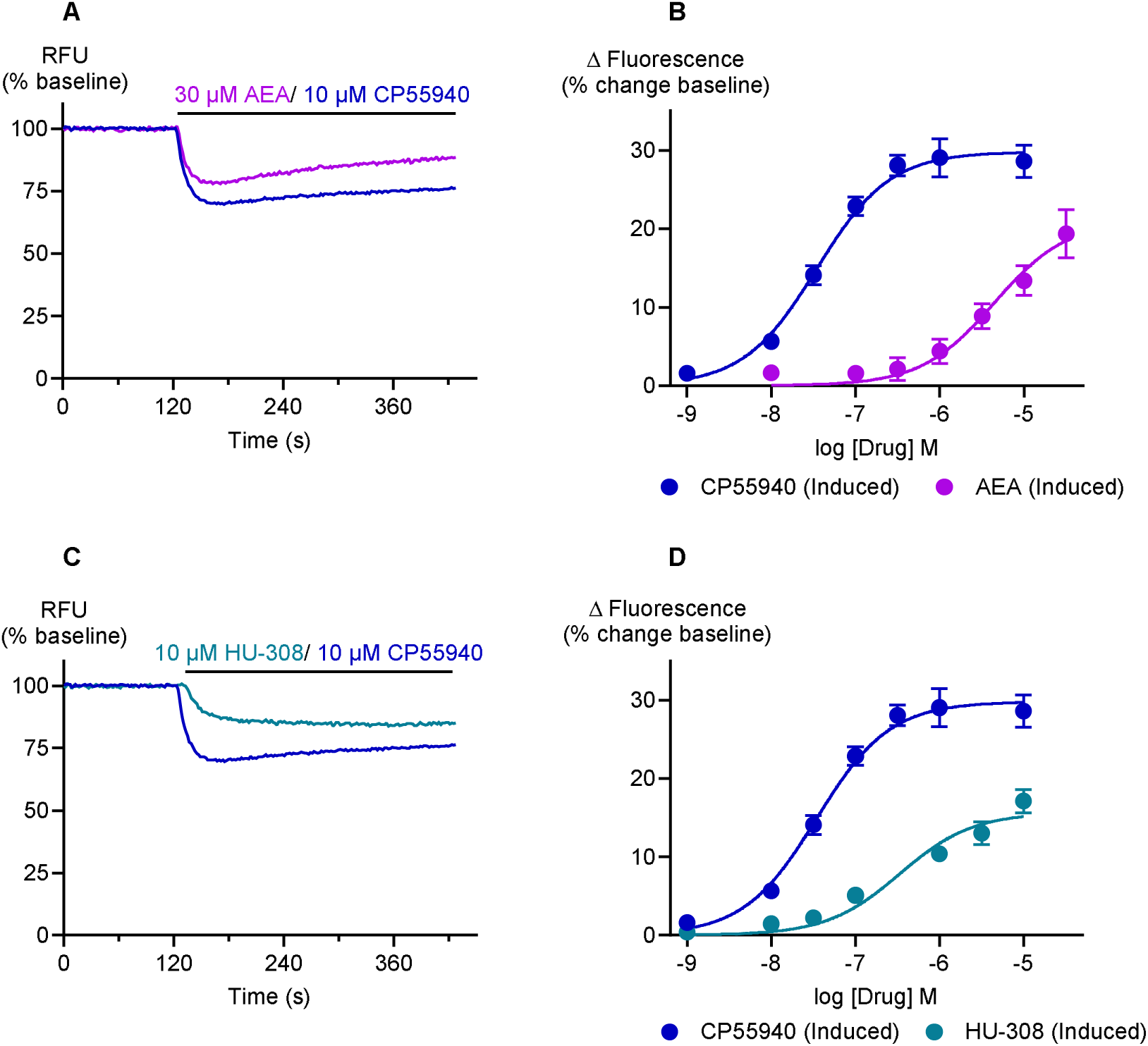
Signalling of low-efficacy CB_2_ agonists compared to reference agonist, CP55940 in AtT20Flp-InT-REx-CB_2_ cells incubated with tetracycline, in FLIPR assay.

**Table 1:** Operational efficacy (τ) and functional affinity (pK_A_) of tested CB_2_ agonists.

| CB<sub>2</sub><br>agonists | Operational Efficacy ($\tau$) | | Functional<br>affinity<br>$pK_A \pm \text{SEM}$ |
| --- | --- | --- | --- |
| | Uninduced<br>$\pm \text{SEM}$ | Induced<br>$\pm \text{SEM}$ | |
| AK-F-064 | $0.93 \pm 0.05$ | $11.43 \pm 1.7$ | $7.04 \pm 0.08$ |
| CP55940 | $1.15 \pm 0.1$ | $11.00 \pm 0.7$ | $6.53 \pm 0.08$ |
| 2-AG | $1.69 \pm 0.34$ | $10.37 \pm 1.3$ | $5.22 \pm 0.15$ |
| 5F-AB-PICA | $0.67 \pm 0.1$ | $5.94 \pm 0.7$ | $7.11 \pm 0.10$ |
| WIN55212-2 | $0.59 \pm 0.07$ | $5.52 \pm 0.7$ | $7.01 \pm 0.08$ |
| HU-308 | * | $1.41 \pm 0.2$ | $5.12 \pm 0.16$ |
| AEA | * | $1.07 \pm 0.1$ | $5.37 \pm 0.12$ |

\*The responses were undetectable in uninduced cells, so HU-308 and AEA were tested under tetracycline-induced condition only, and the data fit to the “operational model of partial agonists” in GraphPad Prism to determine efficacy (see Methods).

The maximum response of CP55940, obtained in tetracycline-induced cells, was significantly higher relative to uninduced cells (E_max_= 29.0 ± 1.2% and 14.2 ± 1.0%, p < 0.0001; Figure 3A and 3B). This difference in maximum response reaffirms our hypothesis that the system can effectively regulate the receptor expression. The efficacy (τ) of CP55940 in induced and uninduced cells was calculated as 11 ± 0.7 and 1.1 ± 0.1, respectively. This indicates that in uninduced conditions, CP55940 occupied approximately 87% of the available receptors to produce a half-maximal response, whereas in induced conditions, only about 9% receptor occupation is needed for the same response. The functional affinity (pK_A_) of CP55940 was estimated as 6.5 ± 0.08 in this system. AK-F-064 had a similar high E_max_ (29.5 ± 1.8%) to CP55940, with a pK_A_ of 7.04 ± 0.08 (Figure 3C and 3D; Table 1). We sought to compare the efficacy of all the ligands tested under tetracycline-induced conditions. A one-way ANOVA showed a significant difference in the mean efficacy of the 7 tested drugs (p < 0.0001, Supplementary Table 5). AK-F-064, CP55940, and 2-AG showed similar high efficacy, significantly different to that of WIN55212-2 and 5F-AB-PICA (Table 1, Supplementary Table 5). The efficacy of HU-308 (τ = 1.41 ± 0.2) and AEA (τ = 1.07 ± 0.1) were similar and the lowest among the tested agonists (Table 1, Figure 5).

5F-AB-PICA exhibited the highest functional affinity (pK_A_ 7.11± 0.1), while HU-308 had the lowest (pK_A_ =5.12 ± 0.16). Two endocannabinoids, 2-AG and AEA, displayed similar functional affinities for CB_2_ (pK_A_= 5.2 ± 0.1 and 5.4 ± 0.1, respectively).

The raw traces represent the decrease in fluorescence in the FLIPR assay mediated by the highest tested concentration of (A) CP55940 (C) AK-F-064, and (E) 2-AG. The CRCs (B, D, and F) illustrate the maximal and submaximal responses of the tested agonists in AtT20Flp-InT-REx-CB_2_ cells with 0.1 µg/mL tetracycline (induced) and without tetracycline (uninduced) preincubation. CRCs data are shown as mean ± SEM (n ≥7).

The raw traces represent the decrease in fluorescence in the FLIPR assay mediated by the highest tested concentration of (A) WIN55212-2 and (C) 5F-AB-PICA. The CRCs (B and D) illustrate the maximal and submaximal responses of the tested agonists in AtT20Flp-InT-REx-CB_2_ cells with 0.1 µg/mL tetracycline (induced) and without tetracycline (uninduced) preincubation. CRCs data are shown as mean ± SEM (n ≥7).

The raw traces represent the decrease in fluorescence in the FLIPR assay mediated by (A) AEA and CP55940 and (B) HU-308 and CP55940. The CRCs illustrate the maximum responses of (B) AEA and (D) HU-308, in relation to CP55940 in tetracycline-induced AtT20Flp-InT-REx-CB_2_ cells. CRCs data are shown as mean ± SEM (n ≥ 7).

### Evaluation of the initial rate of signal generation as a kinetic measure of ligand efficacy

We also assessed the signal generation rate of the CB_2_ receptor after application of the maximally effective concentration of five agonists in a situation where there were no spare receptors. The response of two drugs, HU-308 and AEA, was too small to be quantified in uninduced cells. The Initial rate (IR_max_) values of five agonists were summarised in Table 2.

**Table 2:**
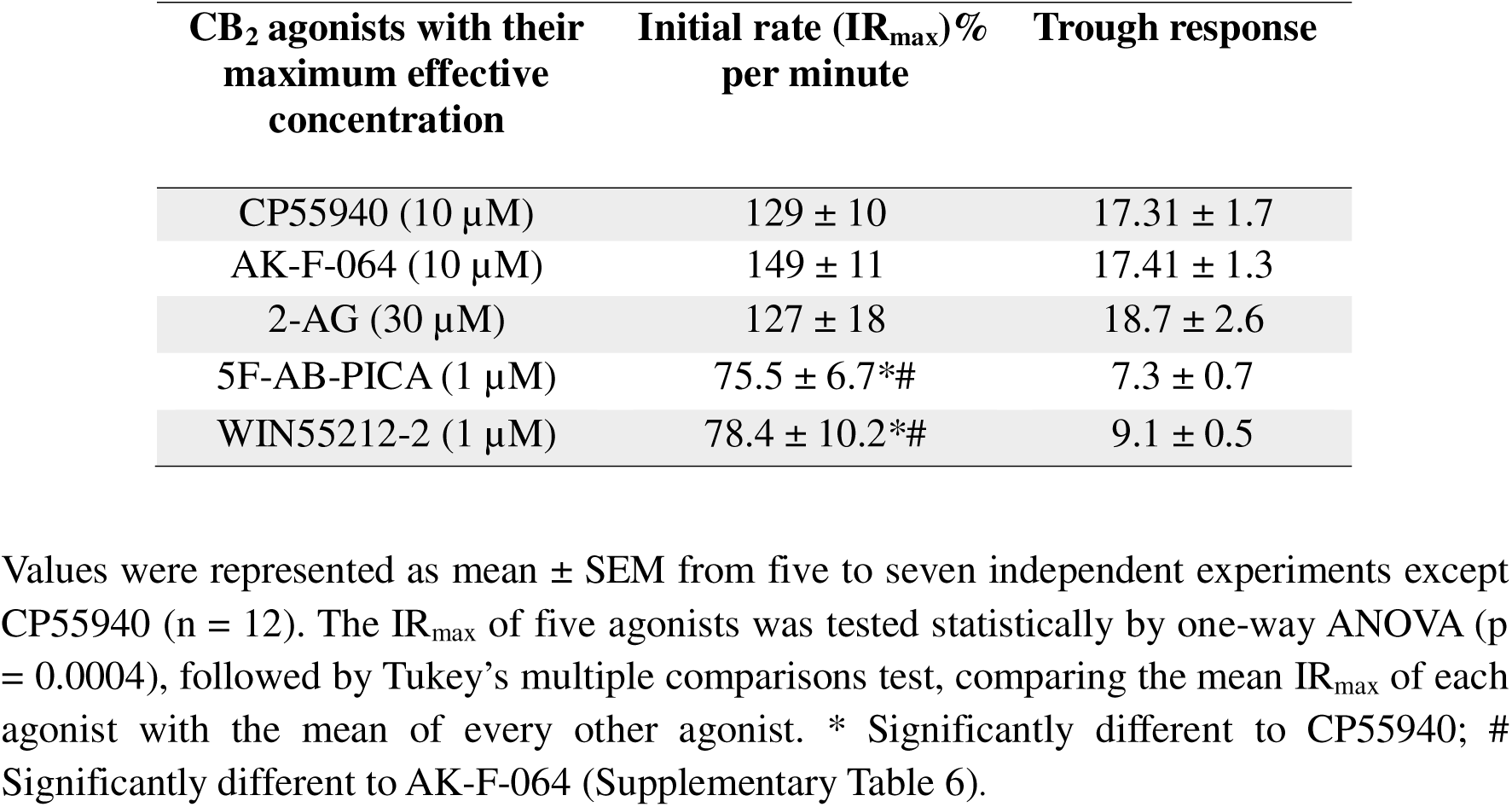
Signal generation rate of CB_2_ agonists upon binding with the CB_2_ receptor.

| **CB<sub>2</sub> agonists with their maximum effective concentration** | **Initial rate (IR<sub>max</sub>)% per minute** | **Trough response** |
| --- | --- | --- |
| CP55940 (10 µM) | 129 ± 10 | 17.31 ± 1.7 |
| AK-F-064 (10 µM) | 149 ± 11 | 17.41 ± 1.3 |
| 2-AG (30 µM) | 127 ± 18 | 18.7 ± 2.6 |
| 5F-AB-PICA (1 µM) | 75.5 ± 6.7*# | 7.3 ± 0.7 |
| WIN55212-2 (1 µM) | 78.4 ± 10.2*# | 9.1 ± 0.5 |

The initial rate of signal generation per minute and the maximum response of CP55940, AK-F-064, and 2-AG were similar. Analysis of the IR_max_ using a one-way ANOVA showed that there was a significant difference among the agonists (p < 0.0004). Subsequent comparisons of the means using Tukey’s multiple comparisons test showed that the IR_max_ for 5F-AB-PICA was significantly different to CP55940 and AK-F-064, as was the IR_max_ for WIN55212-2 (Supplementary Table 6). The amplitude of the maximum responses of CP55940, AK-F-064, and 2-AG compared to baseline was approximately double than 5F-AB-PICA and WIN55212-2.

## Discussion

The primary finding of our study was that the use of an inducible receptor expression system (T-REx) in cells natively expressing a G protein-gated K^+^ channel enabled the use of an operational model to provide an effective strategy to quantify the efficacy of CB_2_ agonists. AK-F-064, CP55940, and 2-AG were high efficacy agonists, WIN55212-2 and 5F-AB-PICA were moderate efficacy agonists, and HU-308 and AEA had a significantly lower efficacy than the other drugs. Moreover, the rank order of the agonist efficacy using the operational model was consistent with a different method of estimating efficacy – measuring the initial rate of signal generation derived from kinetic analysis of the change in fluorescent signal arising from activation of GIRK.

Furchgott’s irreversible receptor inactivation approach [18] has previously been used to facilitate direct fitting of data into operational model to estimate the affinity and operational efficacy of an agonist. However, its application is restricted to receptors with available irreversible antagonists, including opioid, histamine, and β-adrenergic receptors [18,20,32–38]. Our system represents the first approach to overcome this limitation and quantify the operational efficacy of CB agonists in the absence of spare receptors, using the T-REx system for controlled receptor expression and direct operational model fitting. A similar strategy was recently applied to the oxytocin receptor, where agonist responses were assessed by modulating receptor density [39]. Thus, the combination of T-REx with the Flp-In site and a GIRK reporter allows the creation of cell lines expressing any G_i_/G_o_-coupled receptor, which may be especially useful for those receptors where there is no readily available irreversible antagonist. Our experimental procedure is robust, straightforward, omitting any washing steps, and removing the possibility of unspecific effects of covalent ligands.

The system’s functionality was not compromised due to transfection, as the SST responses observed in AtT20Flp-InT-REx host cells, empty vector, and AtT20Flp-InT-REx-CB_2_ cells were comparable. However, the SST responses were significantly lower in AtT20Flp-InT-REx-CB_2_ cells when induced with 0.1 µg/mL tetracycline compared to the cells without tetracycline (Figure 2C and 2D), suggesting that the overexpression of CB_2_ might affect the signalling or expression of the native somatostatin receptors in AtT20 cells. High levels of CB_2_ expression (7.6 pmol/ mg CB_2_ protein) in AtT20 cells were previously reported to sequester G proteins, thereby abolishing or reducing the native somatostatin receptor-mediated responses; this has also been reported for CB_1_ [40,41]. The leakiness of the T-REx system is quite common, and the basal expression of CB_2_ without tetracycline did not affect our experiment [42]. The responses of CB_2_ agonists varied across different tetracycline concentrations, suggesting that the inducible expression system is sensitive to the level of tetracycline concentration. It has been reported that high concentrations of tetracycline (1 µg/mL) can alter cellular gene expression [43,44]. Therefore, in this study, 0.1 µg/mL tetracycline was used to minimise tetracycline-induced cellular effects, despite CP55940 eliciting comparable E_max_ following induction with both 0.1 and 1 µg/mL tetracycline. The hyperpolarisation of the cells caused by the activation of the GIRK channel was entirely triggered by the activation of the CB_2_ receptor, as no responses were detected when the agonists were tested in EV (Supplementary Figure 3).

The efficacy of CB_2_ agonists has previously been estimated by comparing E_max_ values to those of a reference “full agonist”, which was used to define a 100% response within the same receptor and assay system [45–47]. For example, 5F-AB-PICA was previously reported as a high-efficacy agonist in the GIRK/MPA assay by normalising its response to 1 µM of CP55940, while in our study, 5F-AB-PICA was twofold less efficacious than CP55940 [47]. The earlier study used a different AtT20Flp-In cell line, where a high, stable level of receptor expression was achieved. Receptor reserve was likely to be significant, and the presence of spare receptors was not considered in these studies [45,48]. These results further illustrate that consideration of E_max_ alone can fail to differentiate among high efficacy agonists, and drugs that elicit similar maximal responses can possess distinct intrinsic efficacies, as the maximal response of the system imposes a ceiling effect [49–51]. Thus, E_max_ cannot provide a quantitative measure of intrinsic efficacy unless there is no upper limit to the maximum response achievable by the system following receptor activation. Agonist efficacy is also influenced by several factors, including cell type, the system where it is measured, receptor reserve, along with effector protein of the system [17,50,52].

CB_2_ agonist-mediated hyperpolarisation of AtT20 cells has been reported to be pertussis toxin-sensitive, indicating involvement of G_i/o_ protein-dependent signalling as well as G_βγ_ subunit-mediated activation of GIRK channels [45,53]. However, activation of the GIRK channels is not a highly amplified process, as it requires four G_βγ_ subunits for complete activation of the GIRK tetramer [54,55]. Low-efficacy agonists may not produce sufficient G_βγ_ release to robustly activate GIRK signalling, which is a potential limitation of this system. G_βγ_-mediated GIRK activation also only reflects one signalling pathway, and it may not reflect efficacy at other signalling pathways such as G_α_ inhibition of adenylyl cyclase or the more complex pathways leading to β-arrestin recruitment. However, the Tet-controlled receptor expression could be used to estimate efficacy at other signalling pathways, including those more effectively activated by low-efficacy agonists.

The initial rate of signalling induced by agonist-bound receptor provides a time-independent measure of agonist efficacy by quantifying receptor signalling prior to the start of regulatory processes such as desensitisation or internalisation [56–58]. It is a robust and system-independent approach that allows for quantification of ligand efficacy in a condition of low receptor expression where all the receptors are saturated. Thus, this method minimises the confounding effects of receptor reserve and signal amplification. The efficacy measures (IR_max_) in the kinetic approach correspond to established efficacy parameters obtained in the operational model (E_max_ and τ), suggesting that the rank order of efficacy should remain consistent across both methods [59]. Therefore, the experimental data generated in our T-REx system were suitable to be analysed by the time-course equation of the kinetic model. The rank order of the efficacy in kinetic terms was quantified as: AK-F-064= CP55940= 2-AG > 5F-AB-PICA= WIN55212-2, which was consistent with the efficacy measured in the T-REx system. This method is suitable as part of experiments to quantify biased agonism, if the decline in fluorescence in the continued presence of agonist can be shown to depend on receptor phosphorylation and recruitment of arrestin [58,59]. Noteworthy, the kinetic data for a very low-efficacy agonist in cells with low receptor conditions may produce a shallow slope that cannot reliably be fitted into the kinetic model.

In conclusion, our study established a framework for estimating agonist efficacy using an inducible receptor expression system. We quantified and ranked the efficacy of CB_2_ agonists using this system. As the development of new CB_2_ synthetic agonists continues to rise, it is important to know the efficacy profile of existing CB_2_ agonists to provide information to assist in the development of drugs with desired clinical outcomes. Overall, our findings illustrate the capability of this innovative system to estimate ligand efficacy using the operational model, which can also be applied to measure the efficacy of other GPCR agonists. This system will be useful to distinguish higher efficacy compounds in the future and allow us to characterise the pharmacological profile of the CB_2_ agonists across different CB_2_ variants.

## Abbreviations Used

2-AG: 2-Arachidonoyl Glycerol (2-AG)
ANOVA: Analysis of variance
ATCC: American Type Culture Collection
AtT20: Mouse pituitary corticotrope tumour
BSA: Bovine Serum Albumin
DMEM: Dulbecco’s modified Eagle’s medium
DMSO: Dimethyl sulfoxide
FBS: Fetal bovine serum
FLIPR: Fluorometric imaging plate reader
FRT: Flp recombinase target
GIRK: G protein-coupled inwardly rectifying potassium
HBSS: Hank’s Balanced Salt Solution
L-15: Leibovitz L-15 medium
MPA: Membrane potential assay
P/S: Penicillin/Streptomycin
PBS: Phosphate-buffered saline
RFU: Relative fluorescence units
SEM: Standard error of the mean
SST: Somatostatin
SSTR: Somatostatin receptor
T-REx: Tetracycline Regulatory Mammalian Expression

## Acknowledgments Authorship confirmation

Tahira Foyzun: Methodology, Investigation, Validation, Formal analysis, Writing-original draft, Mark Connor: Conceptualisation, Review and Editing, Supervision, Marina Junqueira Santiago: Conceptualisation, Methodology, Supervision, Review and Editing, Humayra Zaman: Methodology, Investigation. Michael Kassiou and Annukka Kallinen: Methodology.

## Supplementary Information

**Supplementary Figure 1:**
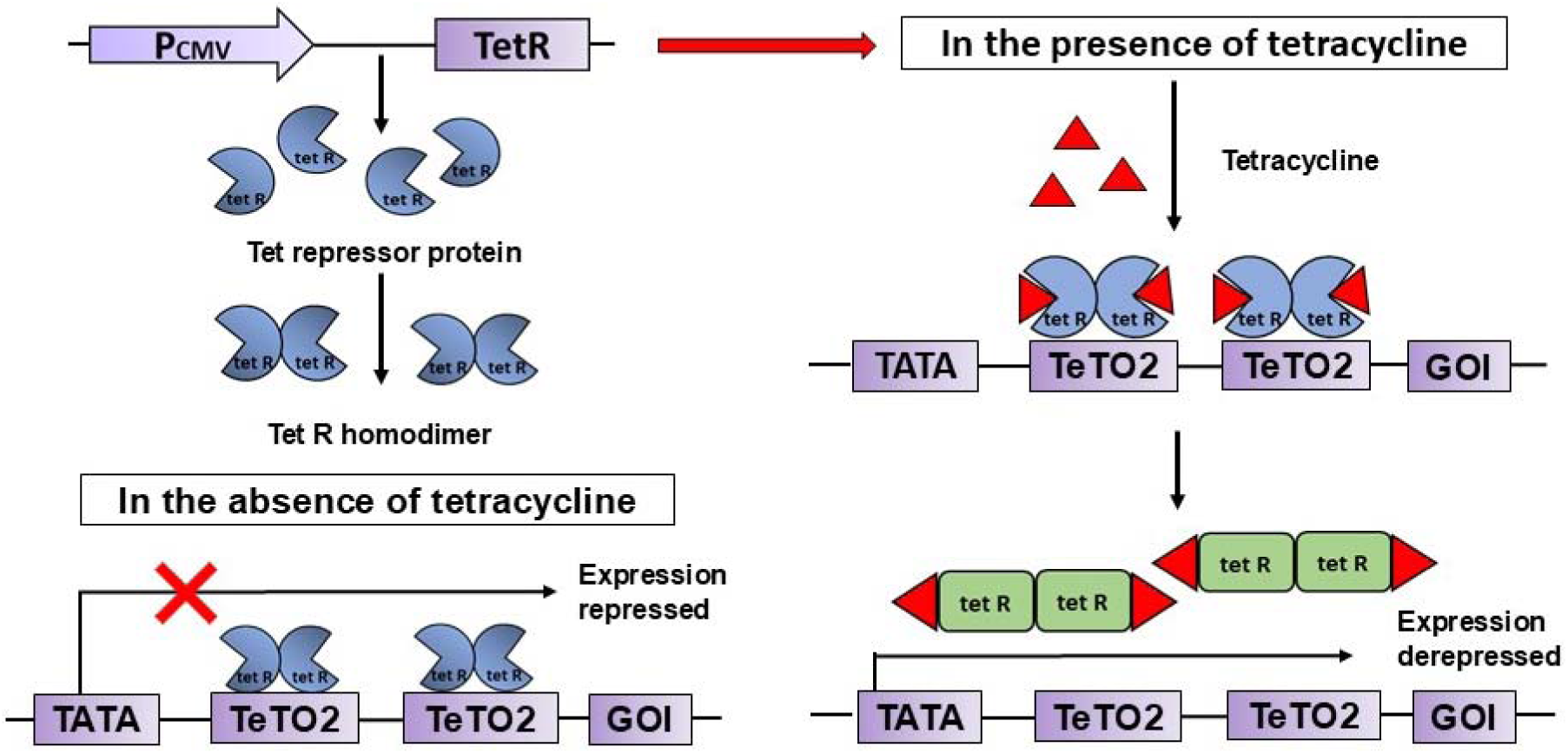
Mechanism of tetracycline induced expression (T-REx) system. The transfected cells constitutively express Tet repressor proteins (TetR) in the culture media and form TetR homodimers. In the absence of tetracycline, the homodimers bind with the tetracycline operator sites (TetO2) positioned between the promoter and the start codon of the gene of interest (GOI), repress the CMV promoter, and prevent transcription and the expression of the GOI. When tetracycline is added to the cells, tetracycline binds with homodimer and forms “Tet homodimer: tetracycline complex” and causes the conformational changes in the homodimer leading to unbinding from the TeTO2 sites. This allows the CMV promoter to activate transcription and express the GOI. (* The image has been recreated from Thermo Fisher Scientific)

**Supplementary Figure 2:**
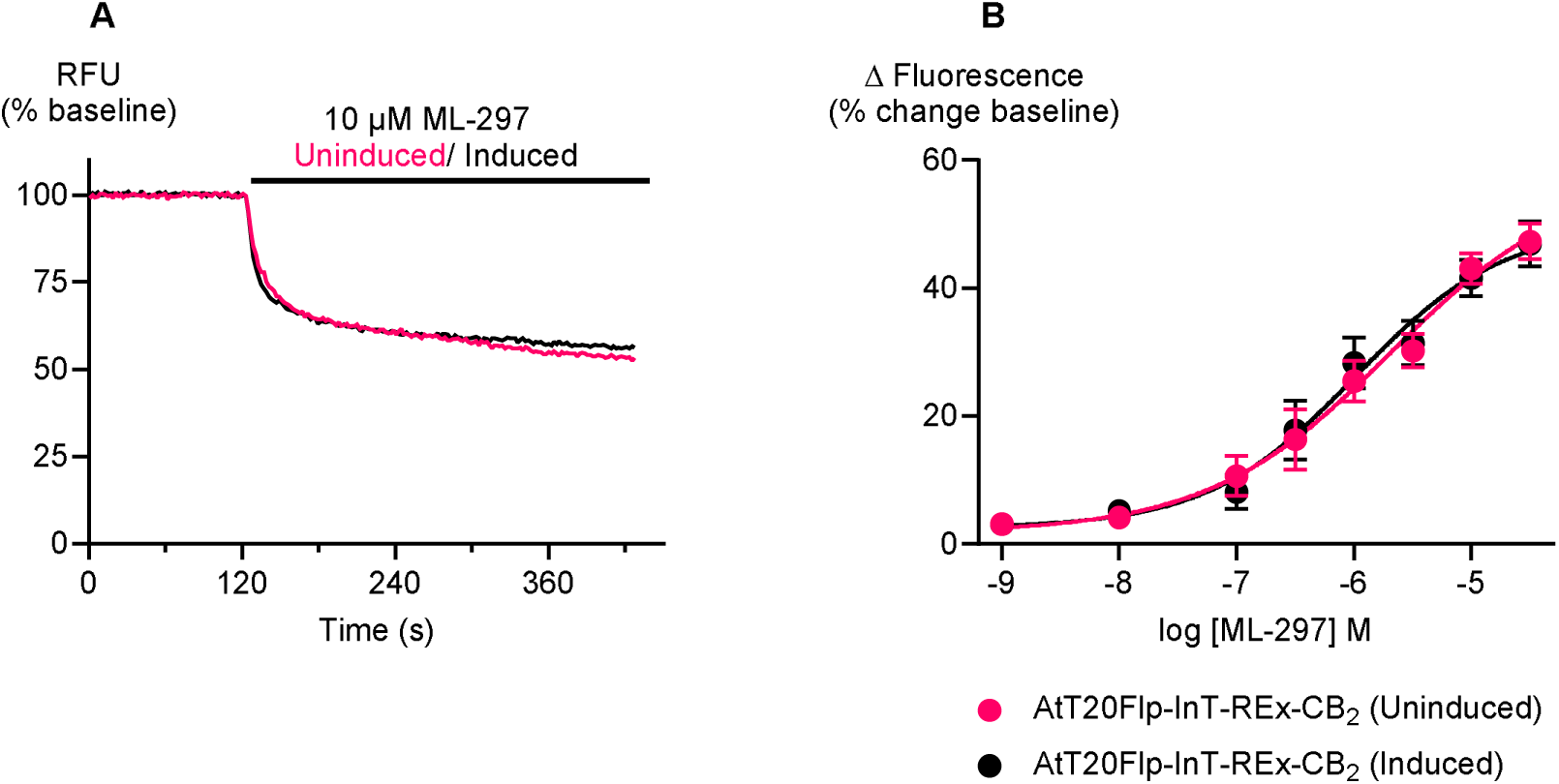
Responses of GIRK channel activator ML-297 in tetracycline-induced and uninduced AtT20Flp-InT-REx-CB_2_ cells. The GIRK channel was activated with ML-297 in tetracycline-induced (0.1 µg/mL) and uninduced cells. (A) Raw traces of the concentration-dependent changes in membrane potential from the ML-297-mediated activation of the GIRK channel, measured by FLIPR assay. (B) Concentration-response curves of ML-297 in tetracycline-induced and uninduced AtT20Flp-InT-REx-CB_2_ cells (n = 5).

**Supplementary Figure 3:**
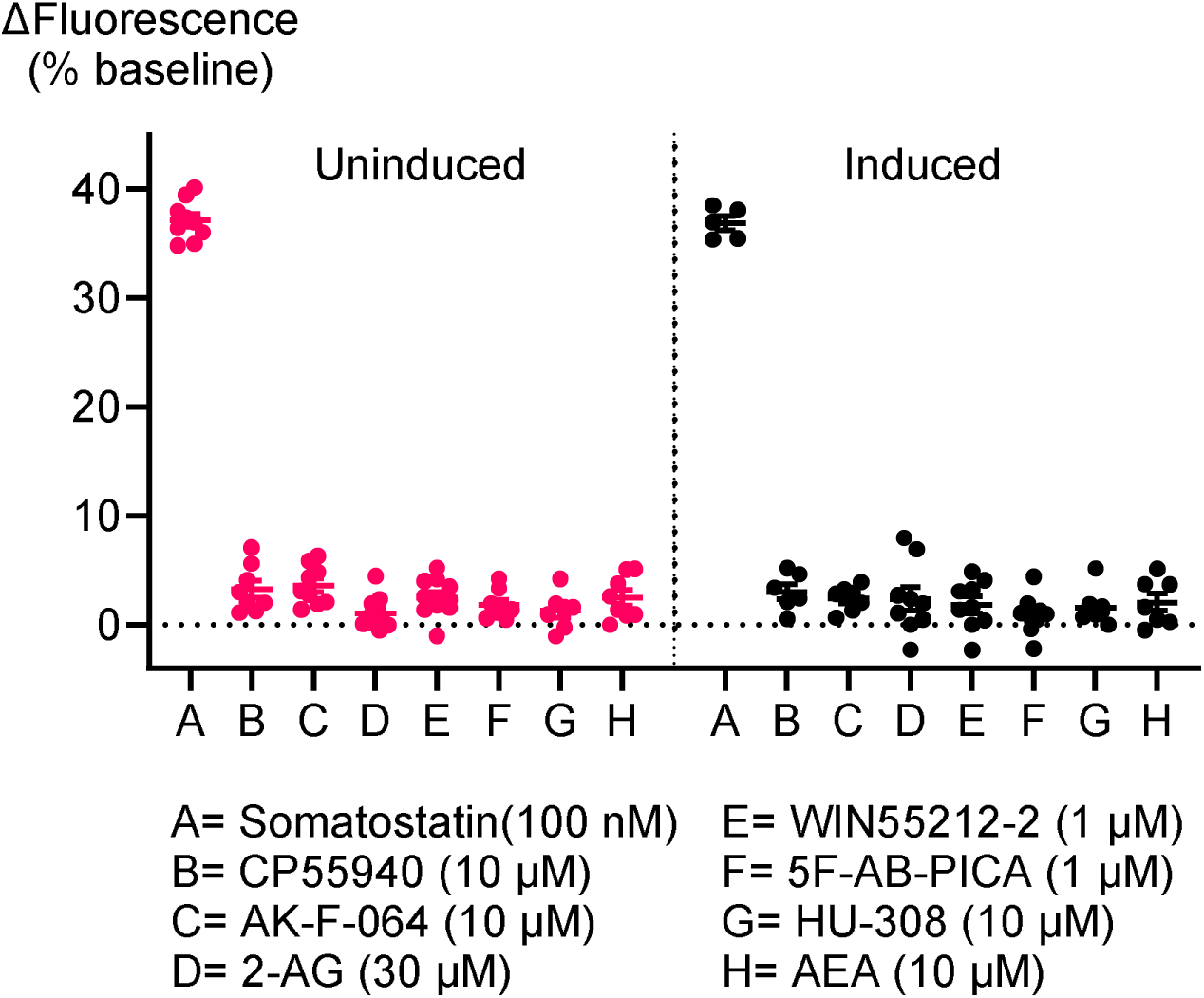
Signalling of somatostatin and tested CB_2_ agonists in empty vector (EV), in the MPA assay. The scatter dot plot represents the changes in fluorescence induced by somatostatin and tested agonists in AtT20Flp-InT-REx-EV cells, incubated with (0.1µg/ mL) and without tetracycline. Somatostatin changed the fluorescence because of the activation of the native somatostatin receptor in EV cells. CB_2_ agonists produced a negligible change in fluorescence because of the absence of CB_2_ receptors. Each dot represents one independent experiment performed in duplicate; bars indicate mean ± SEM (n > 6).

**Supplementary Table 1:**
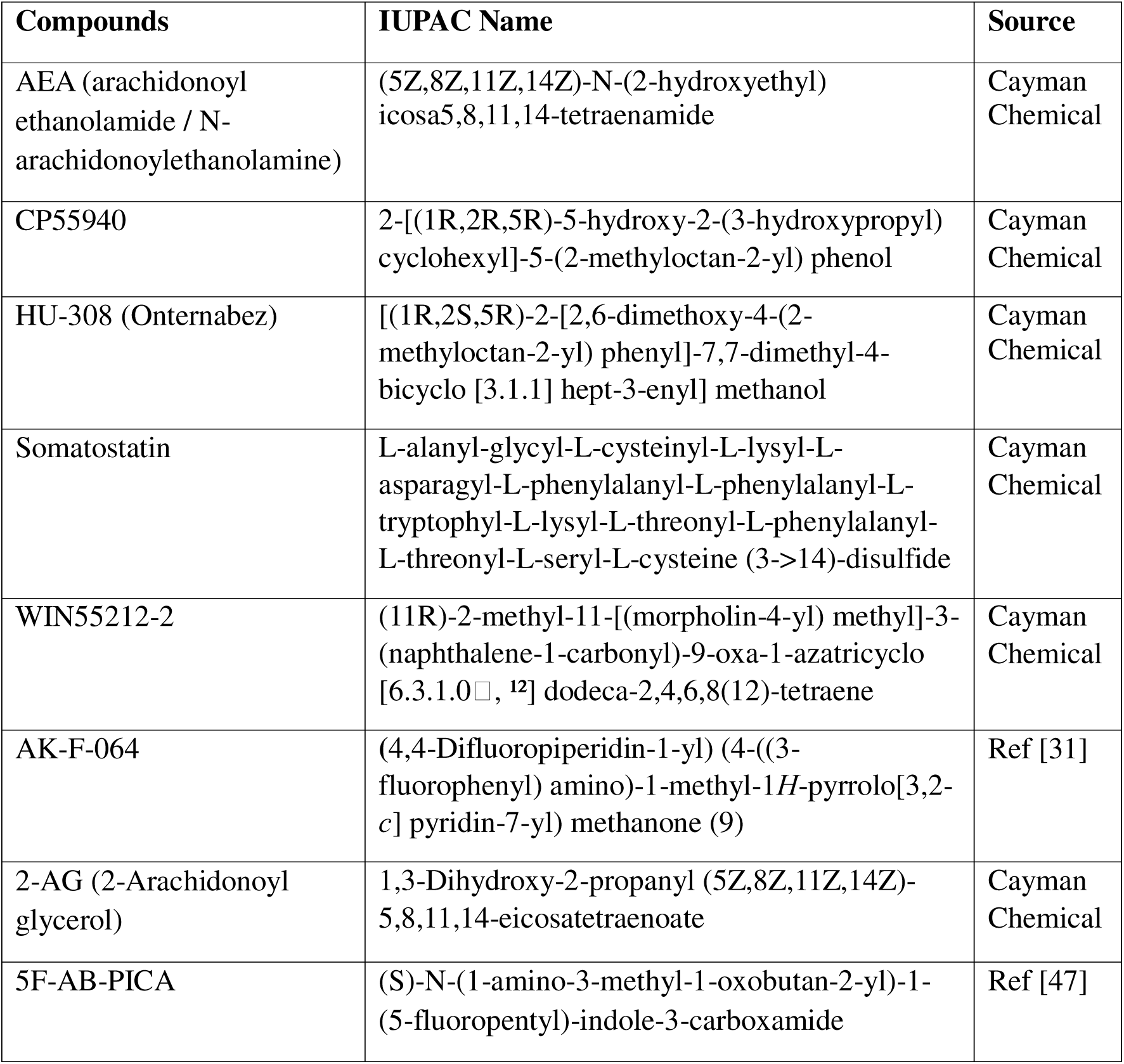
Chemical Nomenclature and sources of tested compounds.

**Supplementary Table 2:**
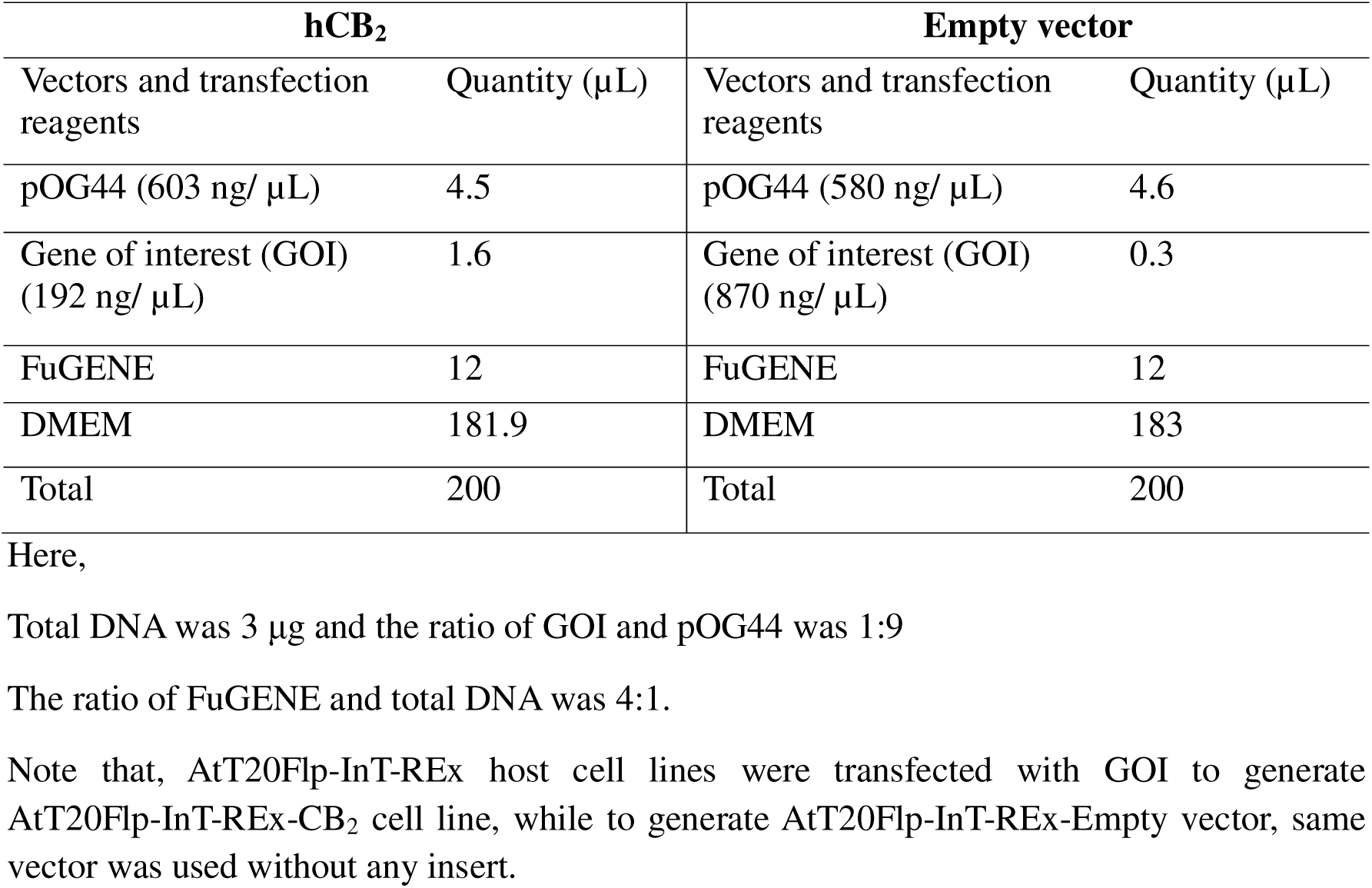
Calculation of transfection reagents.

**Supplementary Table 3:** Composition of low potassium HBSS.

| **Buffer components** | **Concentration (mM)** |
| --- | --- |
| NaCl | 145 |
| HEPES | 22 |
| Na<sub>2</sub>HPO<sub>4</sub> | 0.338 |
| NaHCO<sub>3</sub> | 4.17 |
| KH<sub>2</sub>PO<sub>4</sub> | 0.441 |
| MgSO<sub>4</sub> | 0.407 |
| MgCl<sub>2</sub> | 0.493 |
| Glucose | 5.55 |
| CaCl<sub>2</sub> | 1.26 |

\*pH=7.4 and Osmolarity= 315 ± 15 mOsm/L. The solution was made in milli-Q water and sterilized through 0.22μm filter and stored at 4°C.

**Supplementary Table 4:** The E_max_ and pEC_50_ values of tested CB_2_ agonists in uninduced and induced conditions.

| CB<sub>2</sub> agonists | Uninduced condition |  | Induced condition<br>(induced with 0.1 µg/mL tetracycline) |  |
| --- | --- | --- | --- | --- |
|  | E<sub>max</sub> (%) | pEC<sub>50</sub> | E<sub>max</sub> (%) | pEC<sub>50</sub> |
| **CP55940** | 14.2 ± 1.0 | 7.0 ± .06 | 29.0 ± 1.2 | 7.3 ± 0.06 |
| **AK-F-064** | 13.8 ± 1.9 | 7.3 ± 0.2 | 29.5 ± 1.8 | 8 ± 0.2 |
| **2-AG** | 13.5 ± 2.5 | 5.6 ± 0.1 | 26.5 ± 1.9 | 6 ± 0.1 |
| **5F-AB-PICA** | 10.2 ± 1.0 | 7.6 ± .09 | 25.5 ± 1.4 | 7.8 ± 0.05 |
| **WIN55212-2** | 10.2 ± 0.4 | 7.2 ± .07 | 24.1 ± 1.8 | 7.6 ± 0.05 |
| **HU-308** | * | * | 16.4 ± 1.5 | 6.6 ± 0.2 |
| **AEA** | * | * | 14.2 ± 2.4 | 5.5 ± 0.1 |

\*HU-308 and AEA were tested only in induced conditions

**Supplementary Table 5:**
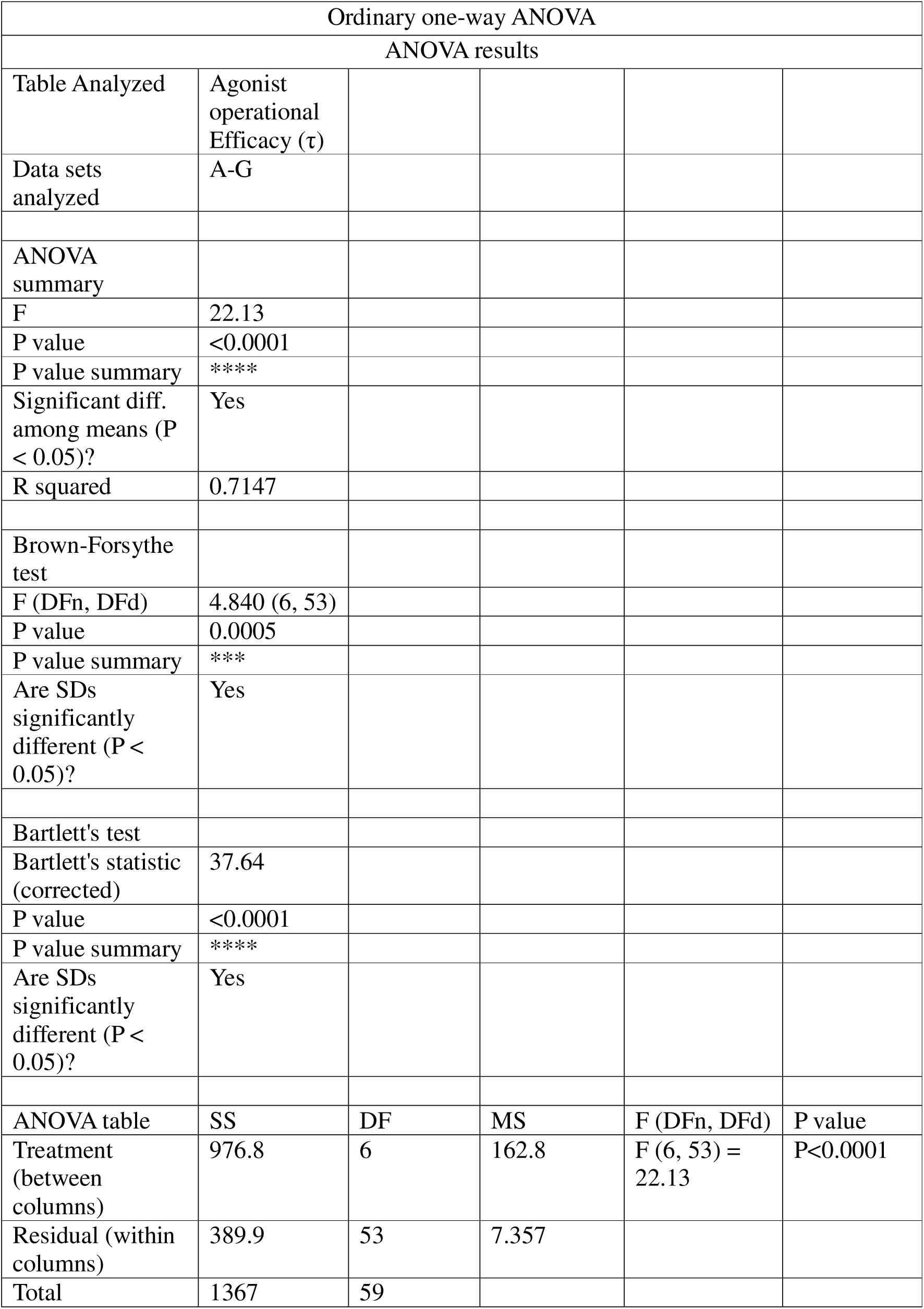
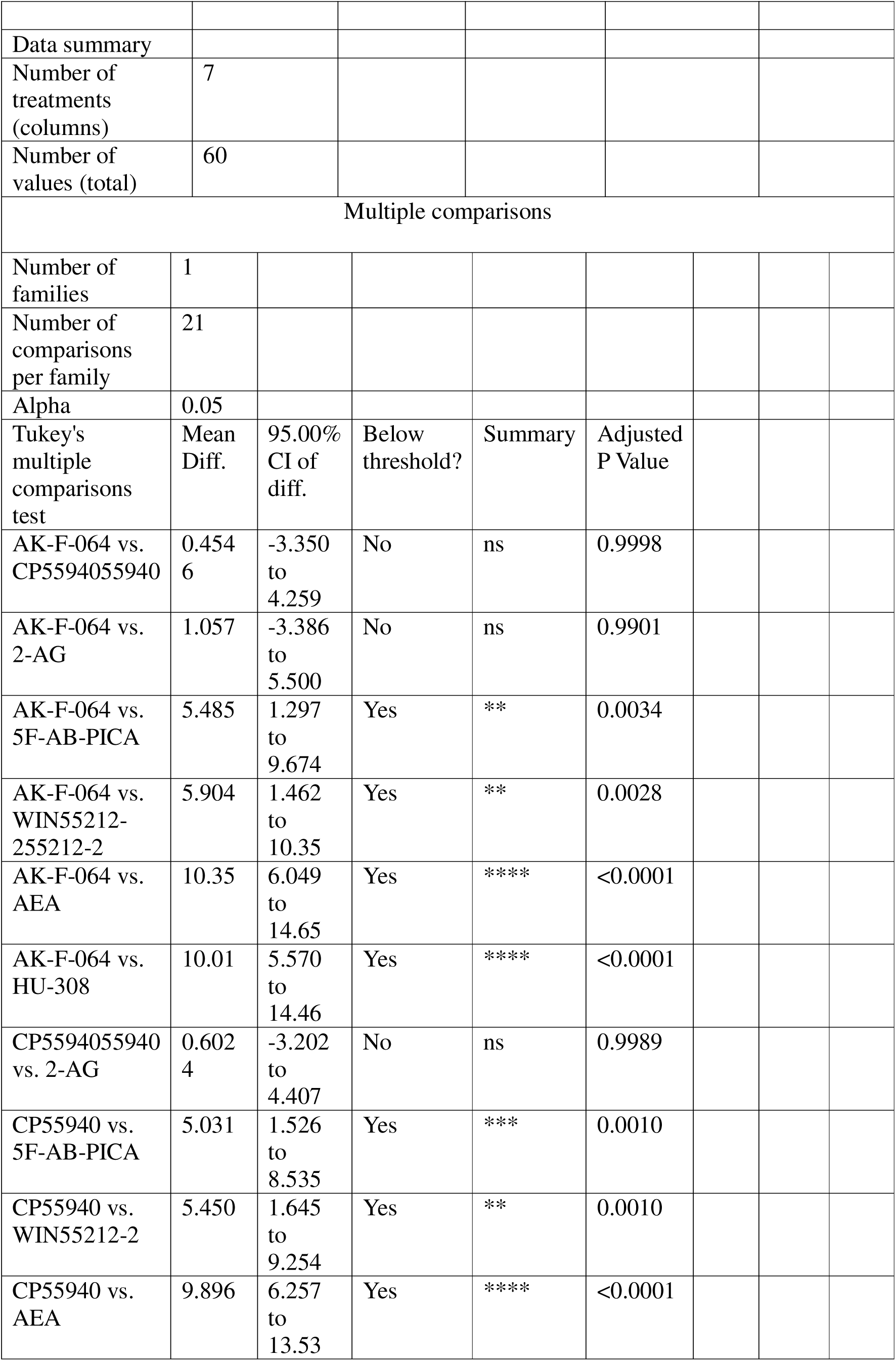
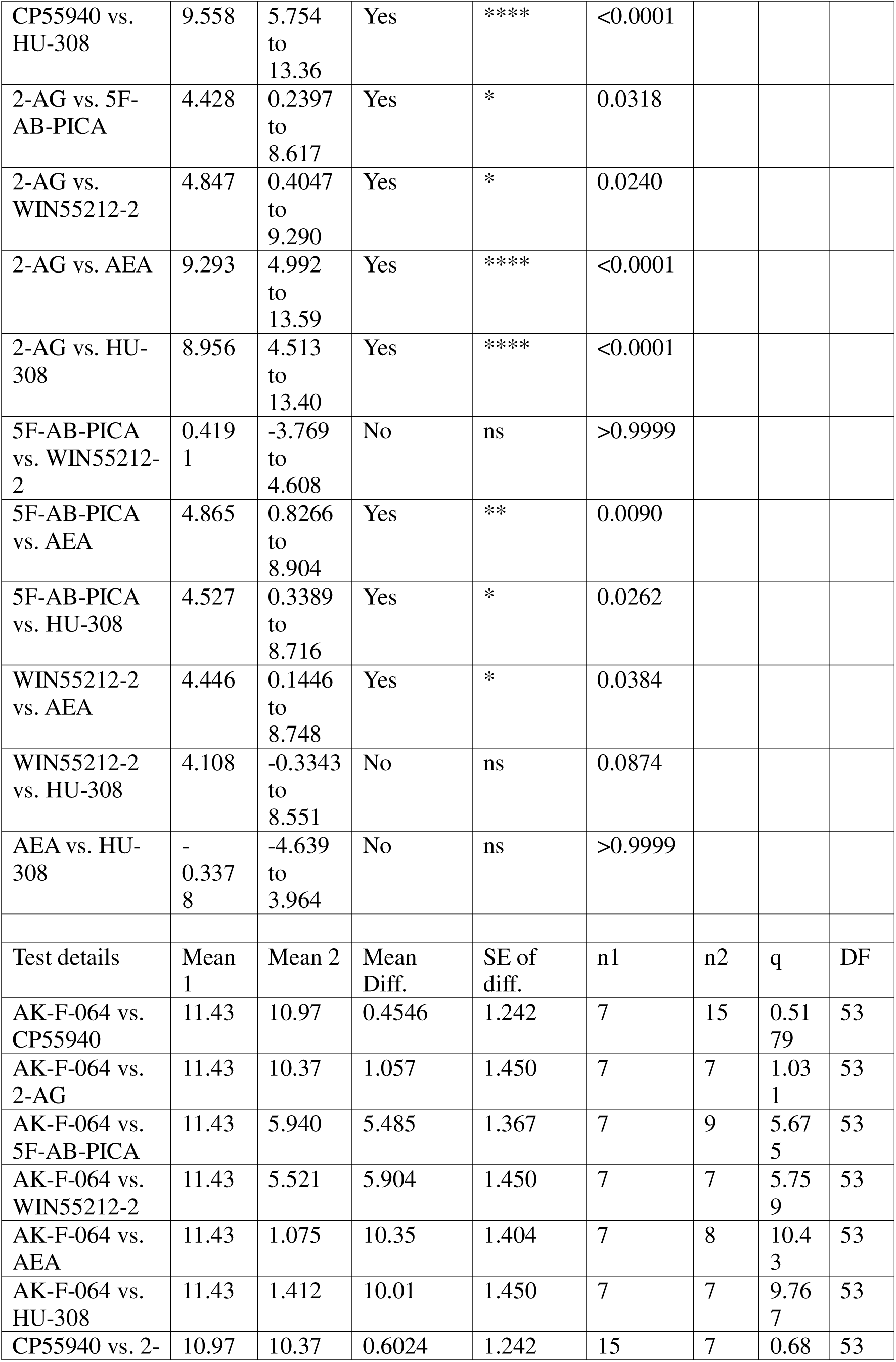
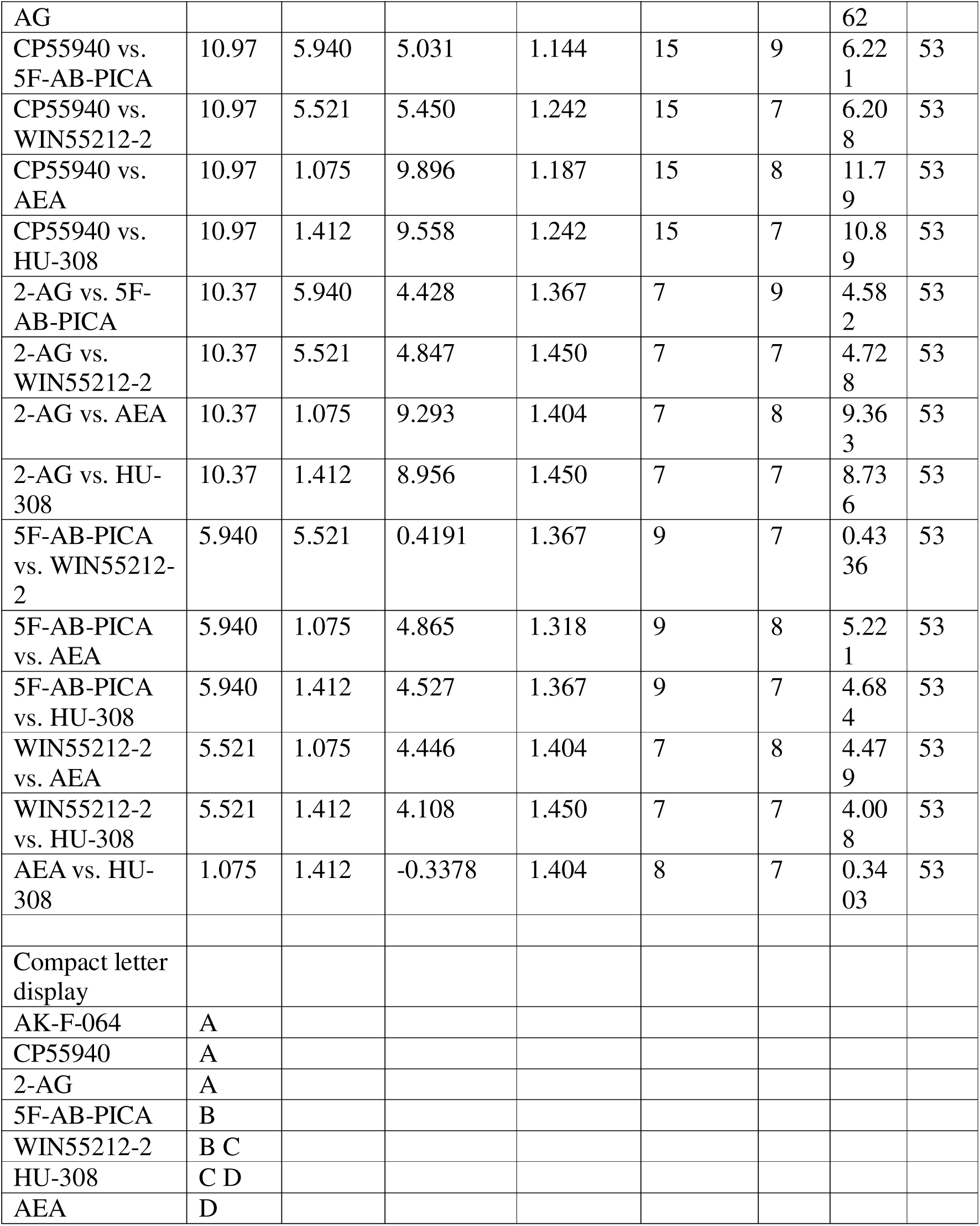
Results of one-way ANOVA and Tukey’s Multiple Comparisons Test for operational efficacy (τ) of the agonists, in conditions with spare receptors (cells were treated with 0.1 µg/mL of tetracycline), obtained from PRISM 10.3.1.

| Ordinary one-way ANOVA |  |  |  |  |  |
| --- | --- | --- | --- | --- | --- |
| ANOVA results |  |  |  |  |  |
| Table Analyzed | Agonist operational Efficacy ($\tau$) | | | | |
| Data sets analyzed | A-G |  |  |  |  |
| ANOVA summary |  |  |  |  |  |
| F | 22.13 |  |  |  |  |
| P value | <0.0001 |  |  |  |  |
| P value summary | **** |  |  |  |  |
| Significant diff. among means (P < 0.05)? | Yes |  |  |  |  |
| R squared | 0.7147 |  |  |  |  |
| Brown-Forsythe test |  |  |  |  |  |
| F (DFn, DFd) | 4.840 (6, 53) |  |  |  |  |
| P value | 0.0005 |  |  |  |  |
| P value summary | *** |  |  |  |  |
| Are SDs significantly different (P < 0.05)? | Yes |  |  |  |  |
| Bartlett's test |  |  |  |  |  |
| Bartlett's statistic (corrected) | 37.64 |  |  |  |  |
| P value | <0.0001 |  |  |  |  |
| P value summary | **** |  |  |  |  |
| Are SDs significantly different (P < 0.05)? | Yes |  |  |  |  |
| ANOVA table | SS | DF | MS | F (DFn, DFd) | P value |
| Treatment (between columns) | 976.8 | 6 | 162.8 | F (6, 53) = 22.13 | P<0.0001 |
| Residual (within columns) | 389.9 | 53 | 7.357 |  |  |
| Total | 1367 | 59 |  |  |  |
| Data summary |  |  |  |  |  |
| Number of treatments (columns) | 7 |  |  |  |  |
| Number of values (total) | 60 |  |  |  |  |
| Multiple comparisons |  |  |  |  |  |
| Number of families | 1 |  |  |  |  |
| Number of comparisons per family | 21 |  |  |  |  |
| Alpha | 0.05 |  |  |  |  |
| Tukey's multiple comparisons test | Mean Diff. | 95.00% CI of diff. | Below threshold? | Summary | Adjusted P Value |
| AK-F-064 vs. CP5594055940 | 0.4546 | -3.350 to 4.259 | No | ns | 0.9998 |
| AK-F-064 vs. 2-AG | 1.057 | -3.386 to 5.500 | No | ns | 0.9901 |
| AK-F-064 vs. 5F-AB-PICA | 5.485 | 1.297 to 9.674 | Yes | ** | 0.0034 |
| AK-F-064 vs. WIN55212-255212-2 | 5.904 | 1.462 to 10.35 | Yes | ** | 0.0028 |
| AK-F-064 vs. AEA | 10.35 | 6.049 to 14.65 | Yes | **** | <0.0001 |
| AK-F-064 vs. HU-308 | 10.01 | 5.570 to 14.46 | Yes | **** | <0.0001 |
| CP5594055940 vs. 2-AG | 0.6024 | -3.202 to 4.407 | No | ns | 0.9989 |
| CP55940 vs. 5F-AB-PICA | 5.031 | 1.526 to 8.535 | Yes | *** | 0.0010 |
| CP55940 vs. WIN55212-2 | 5.450 | 1.645 to 9.254 | Yes | ** | 0.0010 |
| CP55940 vs. AEA | 9.896 | 6.257 to 13.53 | Yes | **** | <0.0001 |

|  |  |  |  |  |  |  |  |  |
| --- | --- | --- | --- | --- | --- | --- | --- | --- |
| CP55940 vs. HU-308 | 9.558 | 5.754 to 13.36 | Yes | **** | <0.0001 |  |  |  |
| 2-AG vs. 5F-AB-PICA | 4.428 | 0.2397 to 8.617 | Yes | * | 0.0318 |  |  |  |
| 2-AG vs. WIN55212-2 | 4.847 | 0.4047 to 9.290 | Yes | * | 0.0240 |  |  |  |
| 2-AG vs. AEA | 9.293 | 4.992 to 13.59 | Yes | **** | <0.0001 |  |  |  |
| 2-AG vs. HU-308 | 8.956 | 4.513 to 13.40 | Yes | **** | <0.0001 |  |  |  |
| 5F-AB-PICA vs. WIN55212-2 | 0.4191 | -3.769 to 4.608 | No | ns | >0.9999 |  |  |  |
| 5F-AB-PICA vs. AEA | 4.865 | 0.8266 to 8.904 | Yes | ** | 0.0090 |  |  |  |
| 5F-AB-PICA vs. HU-308 | 4.527 | 0.3389 to 8.716 | Yes | * | 0.0262 |  |  |  |
| WIN55212-2 vs. AEA | 4.446 | 0.1446 to 8.748 | Yes | * | 0.0384 |  |  |  |
| WIN55212-2 vs. HU-308 | 4.108 | -0.3343 to 8.551 | No | ns | 0.0874 |  |  |  |
| AEA vs. HU-308 | -0.3378 | -4.639 to 3.964 | No | ns | >0.9999 |  |  |  |
| Test details | Mean 1 | Mean 2 | Mean Diff. | SE of diff. | n1 | n2 | q | DF |
| AK-F-064 vs. CP55940 | 11.43 | 10.97 | 0.4546 | 1.242 | 7 | 15 | 0.5179 | 53 |
| AK-F-064 vs. 2-AG | 11.43 | 10.37 | 1.057 | 1.450 | 7 | 7 | 1.031 | 53 |
| AK-F-064 vs. 5F-AB-PICA | 11.43 | 5.940 | 5.485 | 1.367 | 7 | 9 | 5.675 | 53 |
| AK-F-064 vs. WIN55212-2 | 11.43 | 5.521 | 5.904 | 1.450 | 7 | 7 | 5.759 | 53 |
| AK-F-064 vs. AEA | 11.43 | 1.075 | 10.35 | 1.404 | 7 | 8 | 10.43 | 53 |
| AK-F-064 vs. HU-308 | 11.43 | 1.412 | 10.01 | 1.450 | 7 | 7 | 9.767 | 53 |
| CP55940 vs. 2- | 10.97 | 10.37 | 0.6024 | 1.242 | 15 | 7 | 0.68 | 53 |

|  |  |  |  |  |  |  |  |  |
| --- | --- | --- | --- | --- | --- | --- | --- | --- |
| AG |  |  |  |  |  |  | 62 |  |
| CP55940 vs.<br>5F-AB-PICA | 10.97 | 5.940 | 5.031 | 1.144 | 15 | 9 | 6.22<br>1 | 53 |
| CP55940 vs.<br>WIN55212-2 | 10.97 | 5.521 | 5.450 | 1.242 | 15 | 7 | 6.20<br>8 | 53 |
| CP55940 vs.<br>AEA | 10.97 | 1.075 | 9.896 | 1.187 | 15 | 8 | 11.7<br>9 | 53 |
| CP55940 vs.<br>HU-308 | 10.97 | 1.412 | 9.558 | 1.242 | 15 | 7 | 10.8<br>9 | 53 |
| 2-AG vs. 5F-AB-PICA | 10.37 | 5.940 | 4.428 | 1.367 | 7 | 9 | 4.58<br>2 | 53 |
| 2-AG vs.<br>WIN55212-2 | 10.37 | 5.521 | 4.847 | 1.450 | 7 | 7 | 4.72<br>8 | 53 |
| 2-AG vs. AEA | 10.37 | 1.075 | 9.293 | 1.404 | 7 | 8 | 9.36<br>3 | 53 |
| 2-AG vs. HU-308 | 10.37 | 1.412 | 8.956 | 1.450 | 7 | 7 | 8.73<br>6 | 53 |
| 5F-AB-PICA<br>vs. WIN55212-2 | 5.940 | 5.521 | 0.4191 | 1.367 | 9 | 7 | 0.43<br>36 | 53 |
| 5F-AB-PICA<br>vs. AEA | 5.940 | 1.075 | 4.865 | 1.318 | 9 | 8 | 5.22<br>1 | 53 |
| 5F-AB-PICA<br>vs. HU-308 | 5.940 | 1.412 | 4.527 | 1.367 | 9 | 7 | 4.68<br>4 | 53 |
| WIN55212-2<br>vs. AEA | 5.521 | 1.075 | 4.446 | 1.404 | 7 | 8 | 4.47<br>9 | 53 |
| WIN55212-2<br>vs. HU-308 | 5.521 | 1.412 | 4.108 | 1.450 | 7 | 7 | 4.00<br>8 | 53 |
| AEA vs. HU-308 | 1.075 | 1.412 | -0.3378 | 1.404 | 8 | 7 | 0.34<br>03 | 53 |
| Compact letter<br>display |  |  |  |  |  |  |  |  |
| AK-F-064 | A |  |  |  |  |  |  |  |
| CP55940 | A |  |  |  |  |  |  |  |
| 2-AG | A |  |  |  |  |  |  |  |
| 5F-AB-PICA | B |  |  |  |  |  |  |  |
| WIN55212-2 | B C |  |  |  |  |  |  |  |
| HU-308 | C D |  |  |  |  |  |  |  |
| AEA | D |  |  |  |  |  |  |  |

**Supplementary Table 6:**
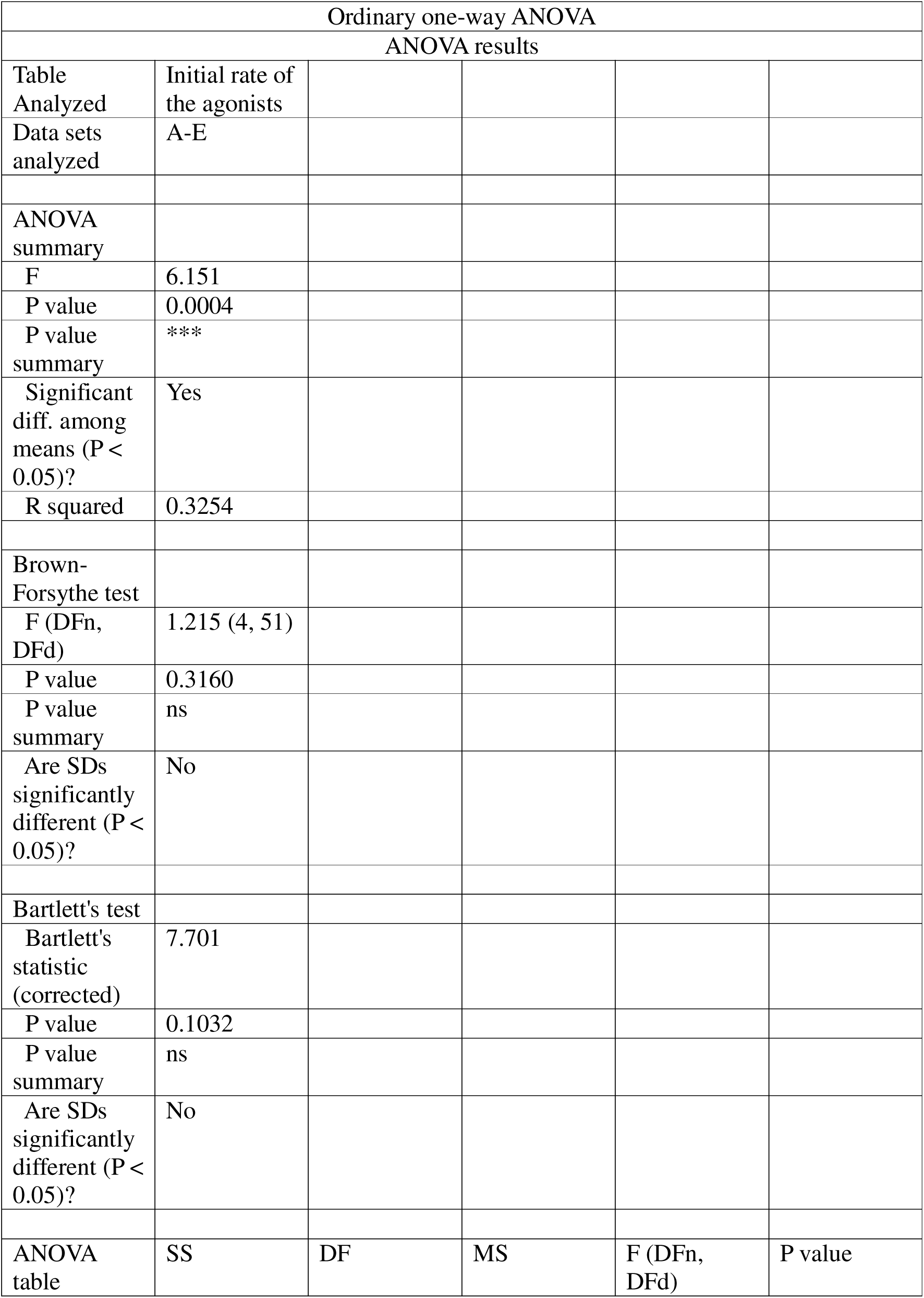
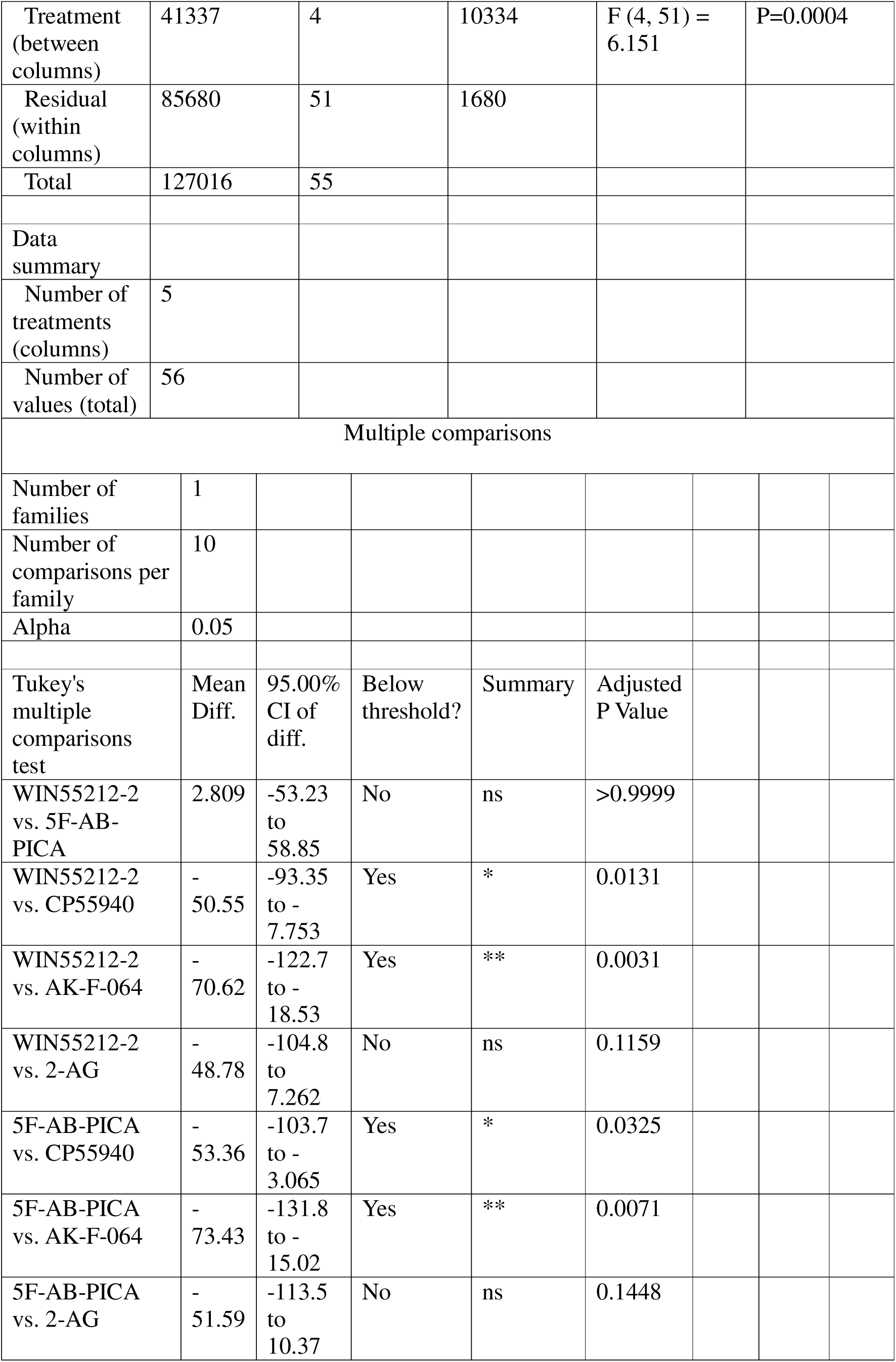
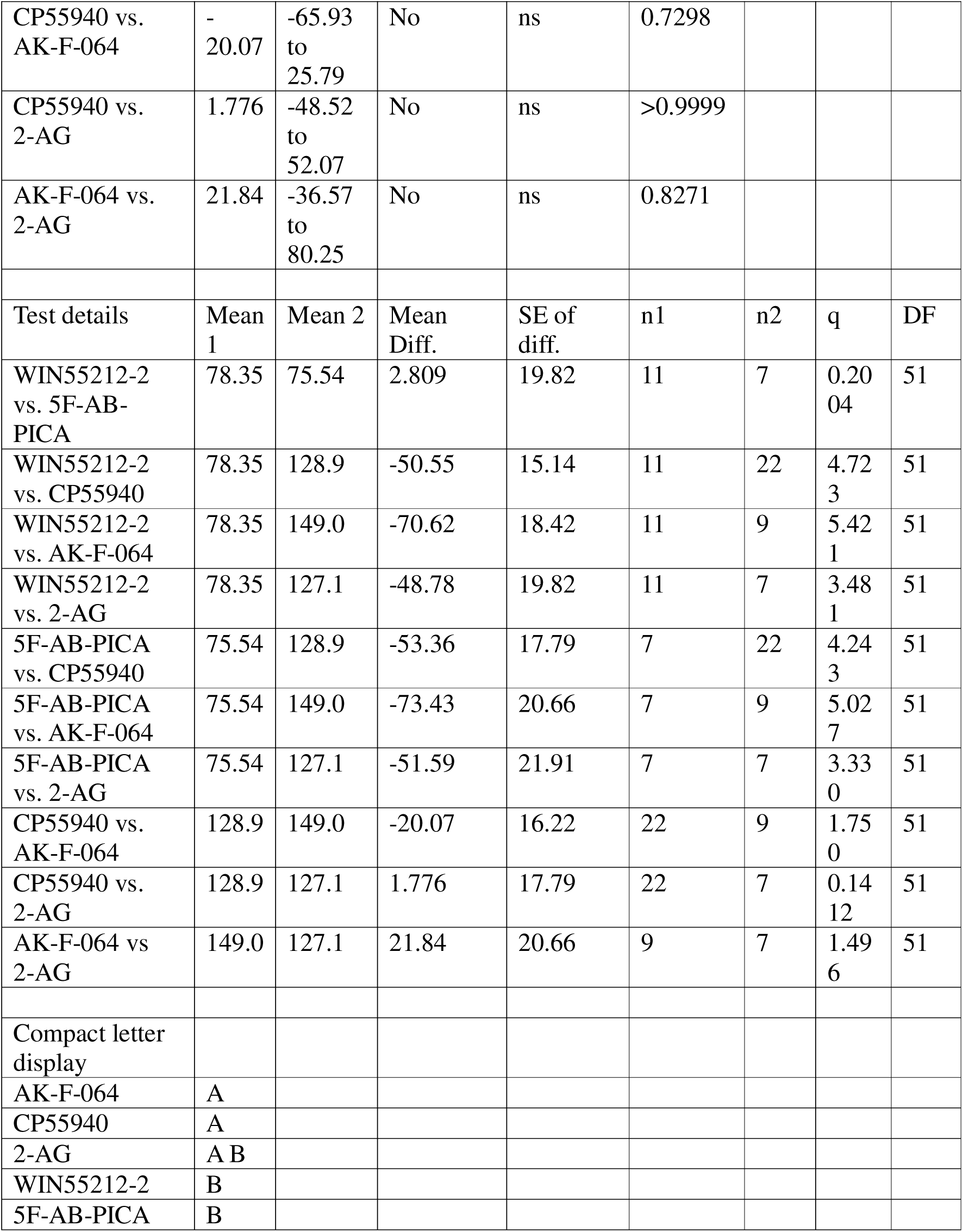
Results of one-way ANOVA and Tukey’s Multiple Comparisons Test for Initial Rate (IR_max_) in conditions with no spare receptors (cells were not induced with tetracycline). From PRISM 10.3.1.

| Ordinary one-way ANOVA |  |  |  |  |  |
| --- | --- | --- | --- | --- | --- |
| ANOVA results |  |  |  |  |  |
| Table Analyzed | Initial rate of the agonists |  |  |  |  |
| Data sets analyzed | A-E |  |  |  |  |
| ANOVA summary |  |  |  |  |  |
| F | 6.151 |  |  |  |  |
| P value | 0.0004 |  |  |  |  |
| P value summary | *** |  |  |  |  |
| Significant diff. among means (P < 0.05)? | Yes |  |  |  |  |
| R squared | 0.3254 |  |  |  |  |
| Brown-Forsythe test |  |  |  |  |  |
| F (DFn, DFd) | 1.215 (4, 51) |  |  |  |  |
| P value | 0.3160 |  |  |  |  |
| P value summary | ns |  |  |  |  |
| Are SDs significantly different (P < 0.05)? | No |  |  |  |  |
| Bartlett's test |  |  |  |  |  |
| Bartlett's statistic (corrected) | 7.701 |  |  |  |  |
| P value | 0.1032 |  |  |  |  |
| P value summary | ns |  |  |  |  |
| Are SDs significantly different (P < 0.05)? | No |  |  |  |  |
| ANOVA table | SS | DF | MS | F (DFn, DFd) | P value |
| Treatment (between columns) | 41337 | 4 | 10334 | F (4, 51) = 6.151 | P=0.0004 |
| Residual (within columns) | 85680 | 51 | 1680 |  |  |
| Total | 127016 | 55 |  |  |  |
| Data summary |  |  |  |  |  |
| Number of treatments (columns) | 5 |  |  |  |  |
| Number of values (total) | 56 |  |  |  |  |
| Multiple comparisons |  |  |  |  |  |
| Number of families | 1 |  |  |  |  |
| Number of comparisons per family | 10 |  |  |  |  |
| Alpha | 0.05 |  |  |  |  |
| Tukey's multiple comparisons test | Mean Diff. | 95.00% CI of diff. | Below threshold? | Summary | Adjusted P Value |
| WIN55212-2 vs. 5F-AB-PICA | 2.809 | -53.23 to 58.85 | No | ns | >0.9999 |
| WIN55212-2 vs. CP55940 | - 50.55 | -93.35 to - 7.753 | Yes | * | 0.0131 |
| WIN55212-2 vs. AK-F-064 | - 70.62 | -122.7 to - 18.53 | Yes | ** | 0.0031 |
| WIN55212-2 vs. 2-AG | - 48.78 | -104.8 to 7.262 | No | ns | 0.1159 |
| 5F-AB-PICA vs. CP55940 | - 53.36 | -103.7 to - 3.065 | Yes | * | 0.0325 |
| 5F-AB-PICA vs. AK-F-064 | - 73.43 | -131.8 to - 15.02 | Yes | ** | 0.0071 |
| 5F-AB-PICA vs. 2-AG | - 51.59 | -113.5 to 10.37 | No | ns | 0.1448 |

|  |  |  |  |  |  |  |  |  |
| --- | --- | --- | --- | --- | --- | --- | --- | --- |
| CP55940 vs.<br>AK-F-064 | -<br>20.07 | -65.93<br>to<br>25.79 | No | ns | 0.7298 |  |  |  |
| CP55940 vs.<br>2-AG | 1.776 | -48.52<br>to<br>52.07 | No | ns | >0.9999 |  |  |  |
| AK-F-064 vs.<br>2-AG | 21.84 | -36.57<br>to<br>80.25 | No | ns | 0.8271 |  |  |  |
| Test details | Mean<br>1 | Mean 2 | Mean<br>Diff. | SE of<br>diff. | n1 | n2 | q | DF |
| WIN55212-2<br>vs. 5F-AB-PICA | 78.35 | 75.54 | 2.809 | 19.82 | 11 | 7 | 0.20<br>04 | 51 |
| WIN55212-2<br>vs. CP55940 | 78.35 | 128.9 | -50.55 | 15.14 | 11 | 22 | 4.72<br>3 | 51 |
| WIN55212-2<br>vs. AK-F-064 | 78.35 | 149.0 | -70.62 | 18.42 | 11 | 9 | 5.42<br>1 | 51 |
| WIN55212-2<br>vs. 2-AG | 78.35 | 127.1 | -48.78 | 19.82 | 11 | 7 | 3.48<br>1 | 51 |
| 5F-AB-PICA<br>vs. CP55940 | 75.54 | 128.9 | -53.36 | 17.79 | 7 | 22 | 4.24<br>3 | 51 |
| 5F-AB-PICA<br>vs. AK-F-064 | 75.54 | 149.0 | -73.43 | 20.66 | 7 | 9 | 5.02<br>7 | 51 |
| 5F-AB-PICA<br>vs. 2-AG | 75.54 | 127.1 | -51.59 | 21.91 | 7 | 7 | 3.33<br>0 | 51 |
| CP55940 vs.<br>AK-F-064 | 128.9 | 149.0 | -20.07 | 16.22 | 22 | 9 | 1.75<br>0 | 51 |
| CP55940 vs.<br>2-AG | 128.9 | 127.1 | 1.776 | 17.79 | 22 | 7 | 0.14<br>12 | 51 |
| AK-F-064 vs<br>2-AG | 149.0 | 127.1 | 21.84 | 20.66 | 9 | 7 | 1.49<br>6 | 51 |
| Compact letter<br>display |  |  |  |  |  |  |  |  |
| AK-F-064 | A |  |  |  |  |  |  |  |
| CP55940 | A |  |  |  |  |  |  |  |
| 2-AG | A B |  |  |  |  |  |  |  |
| WIN55212-2 | B |  |  |  |  |  |  |  |
| 5F-AB-PICA | B |  |  |  |  |  |  |  |

